# Recent reconfiguration of an ancient developmental gene regulatory network in *Heliocidaris* sea urchins

**DOI:** 10.1101/2022.03.03.482868

**Authors:** Phillip L Davidson, Haobing Guo, Jane S Swart, Abdull J Massri, Allison Edgar, Lingyu Wang, Alejandro Berrio, Hannah R Devens, Demian Koop, Paula Cisternas, He Zhang, Yaolei Zhang, Maria Byrne, Guangyi Fan, Gregory A Wray

## Abstract

Changes in developmental gene regulatory networks (dGRNs) underlie much of the diversity of life^1^, but the evolutionary mechanisms that operate on interactions with these networks remain poorly understood. Closely related species with extreme phenotypic divergence provide a valuable window into the genetic and molecular basis for changes in dGRNs and their relationship to adaptive changes in organismal traits. Here we analyze genomes, epigenomes, and transcriptomes during early development in two sea urchin species in the genus *Heliocidaris* that exhibit highly divergent life histories and in an outgroup species. Signatures of positive selection and changes in chromatin status within putative gene regulatory elements are both enriched on the branch leading to the derived life history, and particularly so near dGRN genes; in contrast, positive selection within protein-coding regions have at most a modest enrichment in branch and function. Single-cell transcriptomes reveal a dramatic delay in cell fate specification in the derived state, which also has far fewer open chromatin regions, especially near dGRN genes with conserved roles in cell fate specification. Experimentally perturbing the function of three key transcription factors reveals profound evolutionary changes in the earliest events that pattern the embryo, disrupting regulatory interactions previously conserved for ∼225 million years. Together, these results demonstrate that natural selection can rapidly reshape developmental gene expression on a broad scale when selective regimes abruptly change and that even highly conserved dGRNs and patterning mechanisms in the early embryo remain evolvable under appropriate ecological circumstances.

## MAIN

The well-defined dGRN of early development in sea urchins^2^ provides a powerful framework for investigating the evolution of embryonic patterning mechanisms. Interactions between genes encoding transcription factors and their target genes within this dGRN are almost completely conserved among species that diverged ∼30-40 million years (my) ago^3,4^, with some interactions conserved for ∼225 my^5^, ∼275 my^6^, or even ∼480 my^7,8^. One possible explanation for this observation is developmental constraints, such that early developmental processes are largely immutable given their critical roles in body plan organization and tissue specification^9,10^. Under this scenario, any change in a critical interaction during early development would have widespread effects on later processes, which would almost always be deleterious. Still, an important confound remains untested: the species with deeply conserved developmental mechanisms all share the same life history mode, involving low maternal provisioning and an extended feeding larval phase. Species with derived life histories involving massive maternal provisioning and highly-abbreviated, nonfeeding pre-metamorphic development have evolved on multiple occasions within sea urchins^11-13^, possibly in response to lower or more unpredictable food availability^14^. These species can reveal how conserved regulatory interactions and patterning mechanisms respond to major shifts in selective regimes.

The Australian sea urchin genus *Heliocidaris* includes two recently diverged species: *H. tuberculata* representing the ancestral life history and *H. erythrogramma* the derived state^15^ (Fig.1a). The shift to nonfeeding development radically alters natural selection on development: with feeding no longer necessary, high mortality rates in the plankton^16^ impose strong selection to decrease time to metamorphosis^12^. Numerous anatomical features and gene expression profiles of early development that are broadly conserved among sea urchins differ markedly between these closely related species^17-20^(Fig.1b). We sought to learn whether these recently evolved differences are merely superficial and mask deeply conserved developmental mechanisms, or whether they are the product of substantive evolutionary changes in early cell fate specification and dGRN organization. Evidence for the former would suggest developmental constraints play an important role in limiting genetic and regulatory composition of the ancestral GRN, whereas support for the latter would point to a flexible morphogenetic system derived from an embryonic program conserved at least in part by stabilizing selection that is adaptable to alternative developmental life histories.

**Figure 1.**
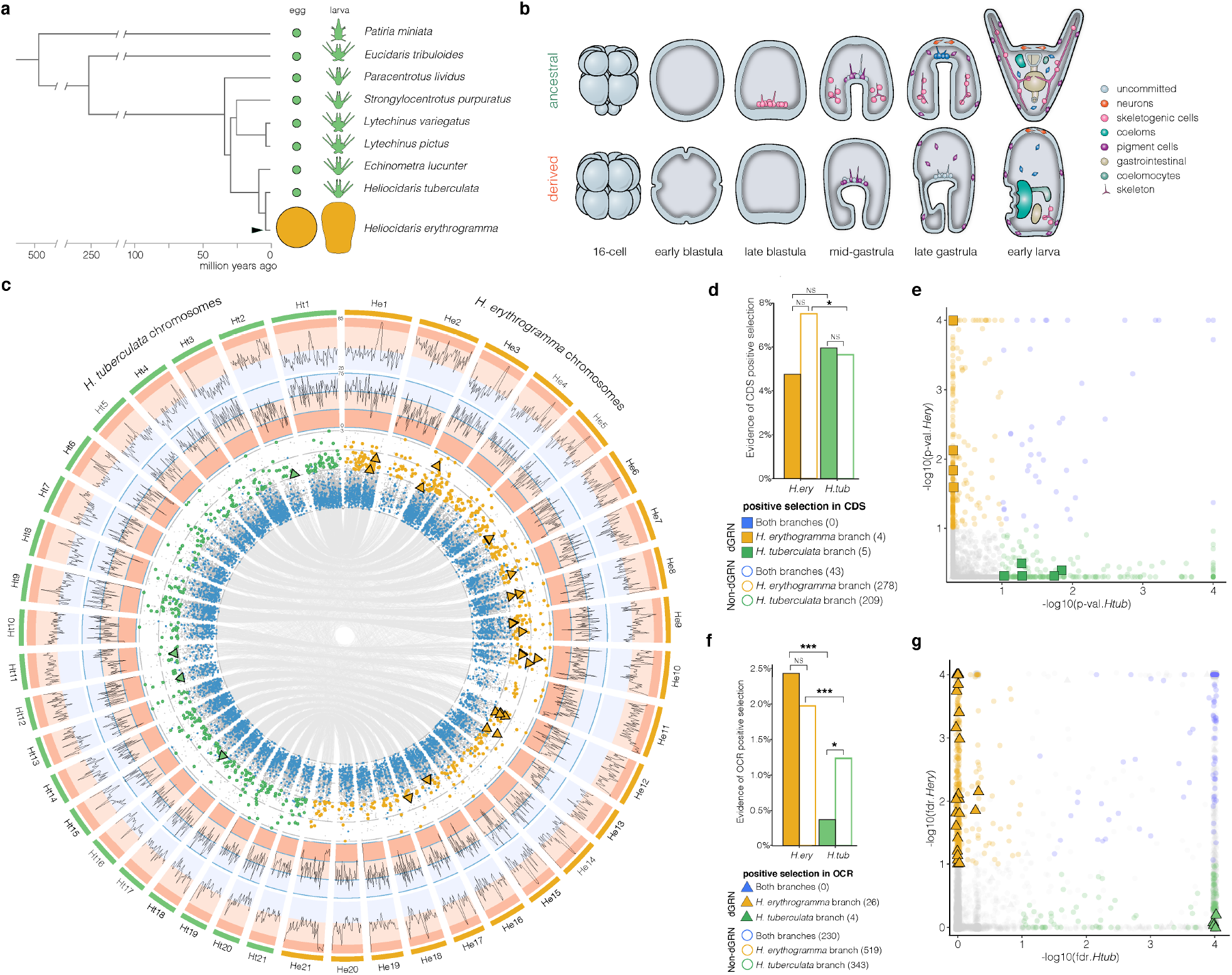
Evolution of life history and genomes. **a**, Feeding larval development (planktotrophy: green) represents the ancient and ancestral life history in sea urchins (seastar *P. miniata* represents the outgroup)^11^. Non-feeding larval development (lecithotrophy: orange) has evolved on multiple occasions, including recently within the genus *Heliocidaris* (arrowhead)^9^. **b**, Evolution of non-feeding development in *H. erythrogramma* (bottom) included dramatic modifications to otherwise broadly conserved developmental mechanisms, including changes in cleavage geometry, cell fate specification, and morphogenesis^16,18^. **c**, Chromosome-scale genome assemblies of *H. tuberculata* (green) and *H. erythrogramma* (orange). Outer ring: repetitive element content; middle ring: gene content, inner ring: zeta values in OCRs from the selection analyses. Colored points indicate statistically elevated zeta values (indicative of positive selection) within a single OCR on the branch leading to *H. tuberculata* (green) or *H. erythrogramma* (orange). Blue points indicate highly conserved OCRs (top 10% of phastCon scores). Triangles denote OCRs with signature of branch-specific positive selection located near dGRN genes. Synteny lines between chromosomes denote locations of 1-1 orthologous OCRs between *Heliocidaris* species. **d**, Signatures of positive selection in protein-coding sequences (CDS) of 84 dGRN and 3,832 non-dGRN single-copy orthologs. Evidence of selection is slightly enriched on the *H. erythrogramma* branch, but dGRN genes show no difference between branches. **e**, P-values of likelihood ratio test for positive selection in CDS on the branch leading to each species (color indicates significant p-values; squares indicate dGRN genes). **f**, Signatures of positive selection within single-copy OCRs near dGRN (n = 1069) and non-dGRN (n = 26,253) genes are overall much higher on the *H. erythrogramma* branch (a single gene can have multiple OCRs associated with it). For OCRs near dGRN genes, this difference is notably amplified: signatures of positive selection are depleted relative to non-dGRN genes on the *H. tuberculata* branch but substantially elevated on the *H. erythrogramma* branch. **g**, P-values of likelihood ratio test for positive selection in OCRs on the branch leading to each species. Fisher exact test, two-sided-*: p-value < 5e-2; ***: p-value < 5e-4. OCR: open chromatin region; dGRN: developmental gene regulatory network; CDS: coding sequence.

## RESULTS

### Natural selection has sculpted the regulatory landscape of *H. erythrogramma*

We took advantage of the recent (∼5 my) divergence between the two *Heliocidaris* species to carry out detailed analyses of orthologous coding and noncoding regions of the genome, focusing on the transcription factors and regulatory elements that constitute the backbone of the dGRN and underlie cell fate specification mechanisms (see Supplementary Data 1 for list of 192 dGRN genes). Genomes of *H. erythrogramma* and *H. tuberculata* were each sequenced, assembled into twenty-one full length chromosomes, and annotated (Fig.1c). Genome sequences were then aligned to one another and to that of *Lytechinus variegatus*^21^, an outgroup representing the ancestral life history condition (Fig.1a).

To understand how natural selection altered the genomes of *Heliocidaris* during the evolution of nonfeeding development, we began by testing for evidence of branch-specific positive selection within single-copy protein coding regions^22^. At a genome-wide scale, we found statistical support for modest enrichment of positive selection along the *H. erythrogramma* branch, but not the *H. tuberculata* branch, when considering the full set of genes (Fisher exact test, two-sided: p<1.33×10^−3^). Of note, coding sequences of dGRN genes showed no enrichment of positive selection on either branch (Fig.1d-e). This result provides little support for the idea that changes in transcription factor structure and function are primarily responsible for the extensive modifications in development and life history in *H. erythrogramma*. Scant evidence of positive selection in the coding sequences of dGRN genes likely reflects pleiotropic constraints imposed by the multiple functions that their encoded transcription factors execute during cell type specification and differentiation.

Therefore, we hypothesized that functional changes in regulatory elements are instead largely responsible for these trait changes. We carried out ATAC-seq on the two *Heliocidaris* species and *L. variegatus* to identify open chromatin regions (OCRs) representing putative regulatory elements in early (hatched) blastula stage embryos, by which time initial cell fates have been specified in the ancestral dGRN. Our analyses are based on all OCRs present in at least one species and located within a genomic region with 1:1:1 orthology among all three species. We then tested for branch-specific positive selection within these OCRs using an approach analogous to that described above for protein-coding regions. At a global scale, these putative enhancer and promoter regions are enriched for evidence of positive selection on the branch leading to *H. erythrogramma* (545) relative to *H. tuberculata* (347) (Fisher exact test, two-sided: p<1.33e-11) (Fig.1f-g). This higher incidence in signatures of positive selection specifically within OCRs on the *H. erythrogramma* branch is indicative of positive selection on regulatory element function that is remarkably widespread within its genome and is consistent with our earlier finding that many expression differences between the two *Heliocidaris* species are genetically based in *cis*^23^.

Strikingly, signals of *H. erythrogramma*-specific positive selection are even more enriched when only considering OCRs near dGRN genes (Fig.1f-g; difference in median zeta: 0.182; Fisher exact test, two-sided: p<5.33e-5). In all, 26 putative regulatory elements located near 23 distinct dGRN genes exhibit evidence of positive selection on the *H. erythrogramma* branch, as opposed to just four on the *H. tuberculata* branch (Fig.1c,g). These 23 genes represent 17.0% of the total within the defined dGRN with a nearby OCR, a marked enrichment compared with the remainder of the genome, where positive selection is detected in OCRs near just 5.7% of genes (Fisher exact test, two-sided: p<4.92e-4: Fig.2c).

### Two distinct regulatory mechanisms underlie divergence in transcriptomes

While the accessibility of most OCRs and expression^19,23^ of most genes are conserved between species (see Fig. 2b and Extended Data Fig. 1 for examples), we observed a striking decrease in chromatin accessibility of many putative regulatory elements throughout the *H. erythrogramma* genome relative to both species representing the ancestral life history (Fig.2a). Of 2,625 orthologous, differentially accessible OCRs between developmental modes, 1,795 sites (68.4%) are significantly less accessible in *H. erythrogramma* (e.g. Fig.2b: *hesC*). Because decreased chromatin accessibility can limit transcription factor access to regulatory elements and because most regulatory interactions in the early sea urchin embryo involve activation of transcription^24^, widespread evolutionary reduction in chromatin accessibility throughout the genome in *H. erythrogramma* embryos suggests an important role for evolutionary changes in chromatin configuration for divergence in gene expression, in this case associated with generally decreased or delayed zygotic transcription for many genes. This interpretation is consistent with indications of a broad delay in embryonic cell fate specification in this species^19,25-29^.

**Figure 2.**
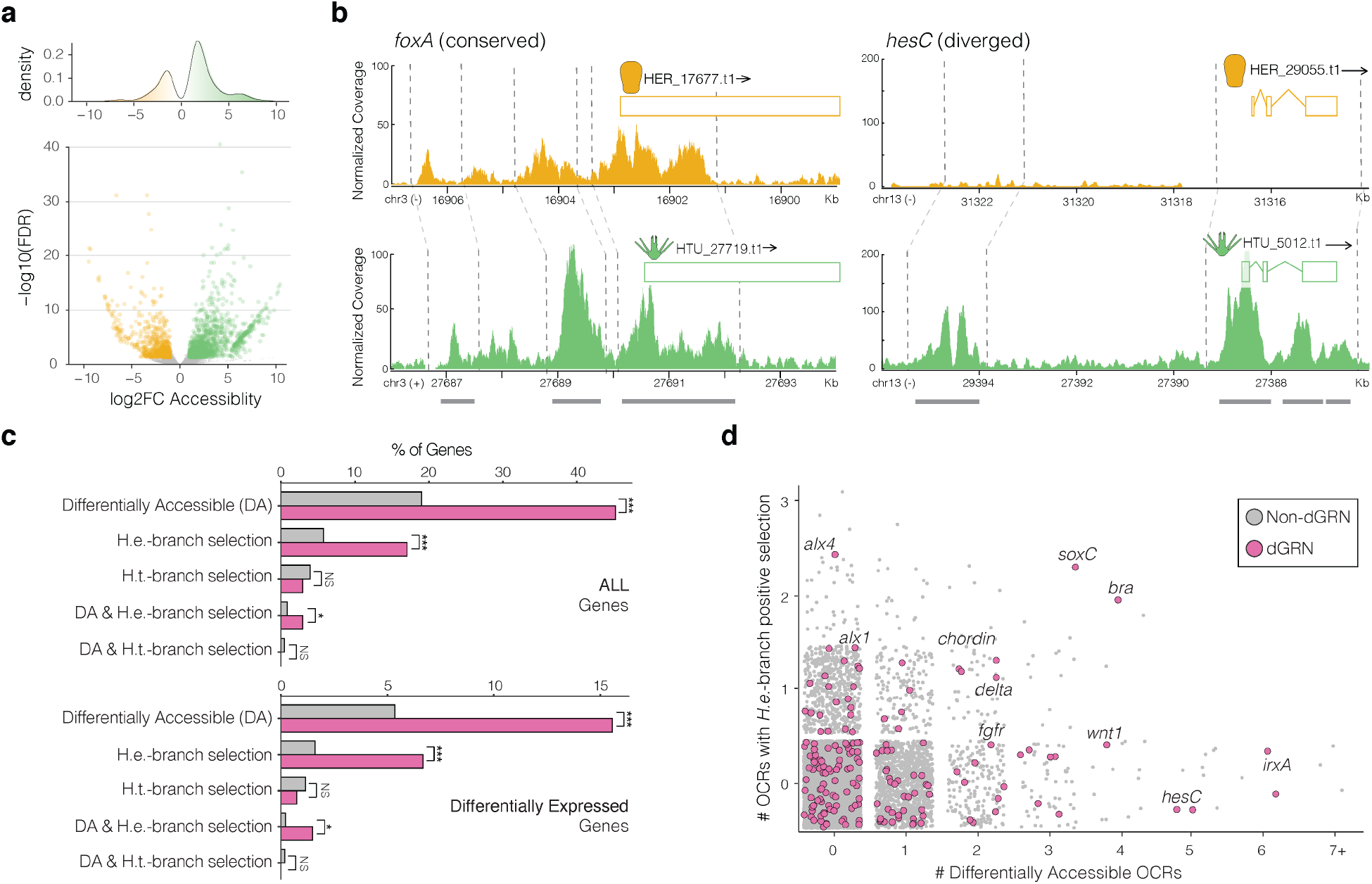
Evolution of open chromatin landscape. **a**, Density and volcano plots of significantly differentially accessible orthologous OCRs between developmental life histories (green = more open in *H. tuberculata* and *L. variegatus*, orange = more open in *H. erythrogramma*). **b**, Examples of conservation (*foxA*) and change (*hesC*) in chromatin accessibility landscape near dGRN genes. See Extended Data Fig. 1 for an additional example of a conserved chromatin landscape nearby *foxN2/3* and corresponding expression domains at blastula stage of both *Heliocidaris* species. **c**, Relationship between chromatin, positive selection, and gene expression. Percentage of all genes (top) and differentially expressed genes (bottom) with at least one OCR nearby that is differentially accessible (DA), has evidence of positive selection on the *H. erythrogramma* (*H.e*.) or *H. tuberculata* (*H.t*.)-branch, or is both DA and has evidence of positive selection on the *H.e*. or *H.t*. branch. **d**, Number of differentially accessible OCRs and OCRs with evidence of positive selection in *H. erythrogramma* for a given gene. Fisher exact test, two-sided-*: p-value < 5e-2; *** p-value < 5e-4. OCR: open chromatin region; dGRN: developmental gene regulatory network.

In a previous study^19^ we analyzed changes in temporal gene expression profiles during early development within *Heliocidaris* and found that the largest changes are concentrated on the branch leading to *H. erythrogramma* and are enriched for developmental regulatory genes generally and dGRN genes specifically. Results reported here suggest that these derived expression profiles are the product of two distinct molecular mechanisms that alter transcription factor binding: changes in nucleotide sequence and changes in chromatin configuration (Fig.2c-d). The former may alter protein:DNA binding while the latter may alter protein access to regulatory elements. Both modes of regulatory evolution are concentrated near dGRN genes relative to the rest of the transcriptome in *H. erythrogramma* (Fig.2c). Notably, accelerated sequence evolution or altered chromatin state (or both) is present in an OCR near a differentially expressed dGRN gene ∼3 times more frequently than the rest of the transcriptome in *H. erythrogramma*, while no such relationship is evident in *H. tuberculata*.

The distribution of both mechanisms of regulatory evolution is highly nonrandom within the genome (enriched near differentially expressed genes and near developmental regulatory genes) and phylogenetically (enriched on the *H. erythrogramma* branch). These departures from the null expectation of random distribution (i.e., resulting from genetic drift) suggest that many of the specific changes are adaptive. Adults of the two *Heliocidaris* species occupy overlapping habitats and ranges^30^, making the suite of derived life history traits that evolved on the *H. erythrogramma* branch the most plausible driver for many of the extensive gene regulatory changes.

### The timing of cell fate specification has changed in *H. erythrogramma*

To understand how changes in the regulation of gene expression in the *H. erythrogramma* embryo influenced developmental mechanisms and life history traits, we leveraged information about the ancestral dGRN to examine embryonic cell fate specification. The earliest zygotic patterning event in the ancestral state involves specification of skeletogenic and germ cell fates following two successive unequal cleavages of vegetal blastomeres^31^. We focus here on the well-characterized skeletogenic cell lineage, which rapidly establishes a distinct transcriptional state^32^ and, within 24 hours after fertilization, undergoes an epithelial-to-mesenchymal transition (EMT), fully differentiates, and begins to synthesize a complex larval endoskeleton (Fig.1b). Specification and maintenance of the skeletogenic cell fate is regulated by interactions between ∼11 transcription factors^33^ (Fig.4a). These developmental events and most of the underlying dGRN interactions are conserved across >225 my of sea urchin evolution^6^.

Morphological development of the skeletogenic cell lineage in *H. erythrogramma* differs in several regards from this ancestral state: cleavage divisions are all equal, no cells undergo EMT before gastrulation, and the larval skeleton is delayed and reduced^20^ (Fig.1b). To understand whether underlying developmental mechanisms are conserved despite these overt morphological differences, we carried out single-cell RNA-sequencing (scRNA-seq) of early blastula stage *H. erythrogramma* embryos and compared the results with our published scRNA-seq data from *L. variegatus*^34^ at the same early blastula stage (prior to EMT). We chose this stage because many major cell fates have been specified by early blastula in the ancestral condition^2,31-33^. This result is clearly reflected in the UMAP of *L. variegatus*, which contains seven cell clusters (Fig. 3a), each expressing a distinct suite of regulatory proteins predicted by the dGRN, with skeletogenic cells exhibiting a particularly disparate transcriptional state (Fig.3c; Supplementary Fig. 1). These indications of early cell fate specification and rapid divergence in transcriptional states are also apparent in scRNA-seq data from *Strongylocentrotus purpuratus*^*35*^, another sea urchin representing the ancestral condition (Fig.1a), suggesting early embryonic regulatory interactions are conserved among planktotrophic species and detectable by scRNA-seq.

**Figure 3.**
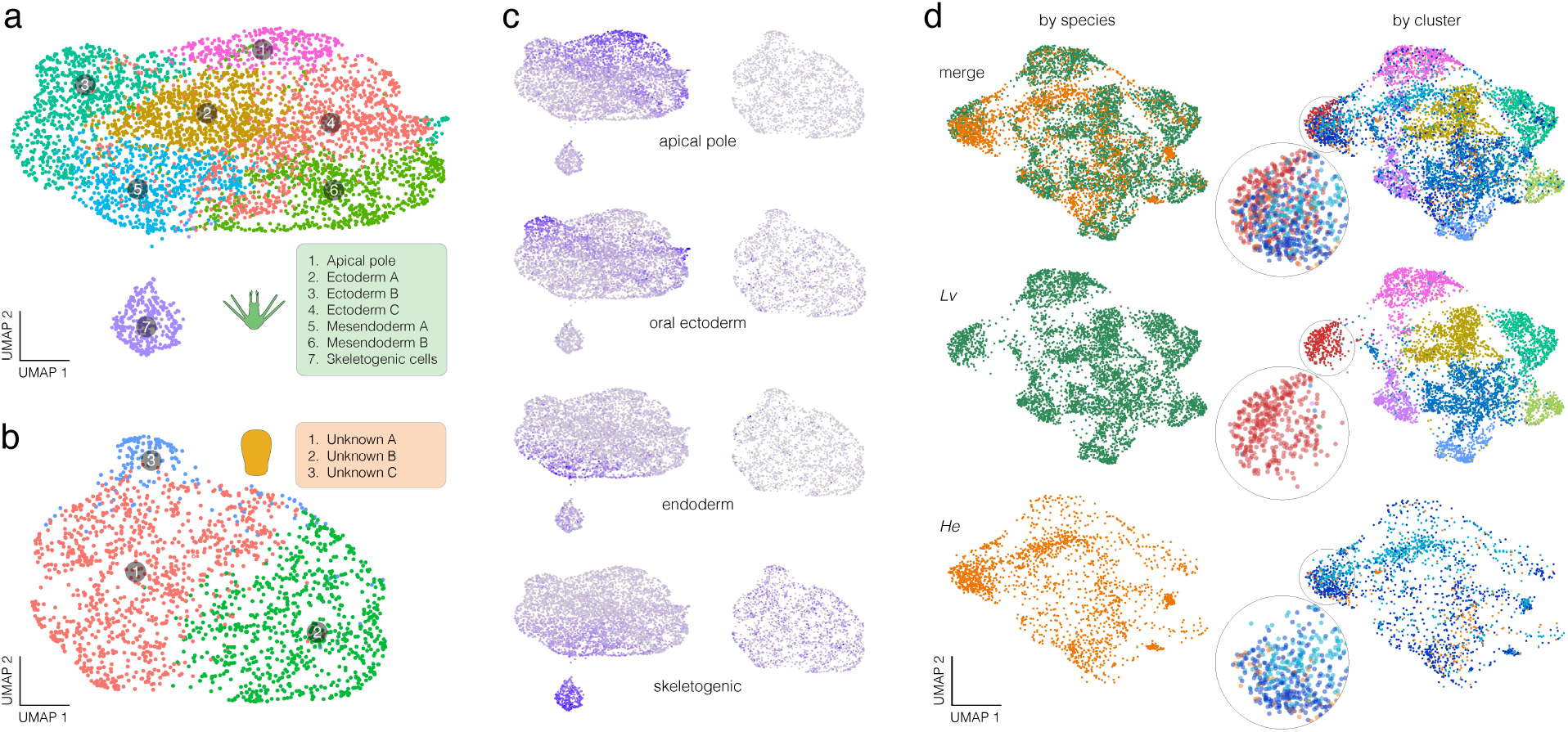
Evolution of transcriptomes. Single cell RNA sequencing of pre-mesenchyme blastula stage embryos from **a**. *L. variegatus* and **b**. *H. erythrogramma* with cells plotted into a UMAP space prior to integration of expression data across species. Colors indicate inferred cell lineages based on expression of marker genes (Supplementary Fig. 1). Note that *H. erythrogramma* shows fewer distinct transcriptional states than *L. variegatus*. **c**, Composite expression scores for four embryonic territories based on combined expression of multiple known marker genes in the ancestral GRN (Supplementary Data 11). These composite expression markers are localized to discrete domains in *L. variegatus* but not in *H. erythrogramma*, suggesting that major embryonic cell lineages have yet to differentiate transcriptionally in this species at this stage of development. **d**, Integrated UMAP of cells from both species, color-coded by species (left) and by cluster (right), and showing both species (top), and separated by species (middle and bottom). *L. variegatus* clusters remain well separated (center right) but *H. erythrogramma* clusters broadly overlap with each other (lower right). As shown in the 2X magnifications, *L. variegatus* clusters contain primarily cells of one color, while cells from all three *H. erythrogramma* clusters are present in appreciable numbers in the same UMAP space (transparency is set to 40% in the insets to circumvent masking; example inset clusters are *L. variegatus* skeletogenic cells, but the same holds for other *L. variegatus* clusters).

In *H. erythrogramma*, only three cell clusters are apparent at the same resolution and developmental stage (Fig.3b). The observation of fewer distinct transcriptional states in *H. erythrogramma* along with less localized expression of known GRN tissue marker genes (Supplementary Fig. 1) suggests a delayed establishment of distinct transcriptional states in the early embryo of this species – a conclusion not attributable technical factors such as analytical parameterization (see Methods), cell number (Extended Data Fig. 2) or genes/UMIs per cell (Supplementary Fig. 2). Given the limited number of cells in this dataset (2065 post-filtering) and representation from a single stage, future work examining later developmental stages will be necessary to fully resolve the timing of cell fate specification in *H. erythrogramma* and associated expression of developmental regulatory factors.

The presence of more numerous transcriptional states in *L. variegatus* is even clearer in an integrated projection of expression for 7,671 one-to-one orthologous genes, where *L. variegatus* clusters remain separated but those of *H. erythrogramma* overlap broadly in UMAP space, suggesting cells of this species have more homogenous transcriptional profiles relative to cells in *L. variegatus* at the same developmental stage (Fig.3d, compare lower center and lower right panels as well as insets). Three independent methods of scRNA-seq data integration are consistent in showing fewer clusters and greater degree of overlap among cells from different clusters in *H. erythrogramma* (Extended Data Fig. 3; see Methods). These findings are consistent with the ATAC-seq results presented above and our earlier lineage-tracing and bulk RNA-seq studies^19, 23, 25, 26^, all of which indirectly point to a delay in fate specification in *H. erythrogramma*.

Further, clusters in the early *H. erythrogramma* do not express similar suites of transcription factors to those in the ancestral state, and none corresponds to the distinctive skeletogenic cell lineage of *L. variegatus* and *S. purpuratus* that is established earlier in their development (Fig. 3c,d; Supplementary Fig. 1,3). For instance, *delta* and *alx1*, which encode critical early regulatory proteins, are expressed exclusively in the skeletogenic cell precursors at the blastula stage in the ancestral state^36,37^. In *H. erythrogramma*, localized transcription of neither *delta* nor *alx1* has commenced (Supplementary Fig. 1). Furthermore, a composite of skeletogenic cell marker gene expression is localized to a discrete cell population in the ancestral state but is not detected in the *H. erythrogramma* embryo (Fig. 3c). Together, these expression differences suggest that the roles of key regulators of the skeletogenic cell fate have evolved during the life history shift.

### The ancient double negative gate within the dGRN is lost in *H. erythrogramma*

To investigate how these roles might differ in the derived developmental mode, we first experimentally perturbed the function of Alx1, which is both necessary and sufficient for skeletogenic cell fate specification in the ancestral state^36^ (Fig.4a). Knocking down Alx1 protein with a translation-blocking morpholino antisense oligonucleotide (MASO) in *H. erythrogramma* eliminates both larval and adult skeleton (Fig.4b; Supplementary Table 1), phenocopying the results of prior experiments in sea urchins representing the ancestral condition^36^. This concordance suggests that the function of Alx1 in skeletogenic cell fate specification is conserved. Zygotic transcription of *alx1*^*19*^ and skeletogenesis^38^ are both markedly delayed in *H. erythrogramma* relative to the ancestral state, but this shift in timing does not by itself indicate a substantive change to the organization of the dGRN.

We next examined HesC, a transcription factor that acts even earlier in the dGRN, repressing transcription of *alx1* outside of the skeletogenic cell lineage (Extended Data Fig. 4). Experimentally eliminating HesC protein in the ancestral dGRN produces a dramatic phenotype, with most cells differentiating as skeletogenic because in the absence of *hesC* repression, *alx1* is broadly transcribed^39^. In *H. erythrogramma*, however, we found that embryos develop normally following HesC knock-down (Extended Data Fig. 4e-f), suggesting that it no longer acts as a repressor of *alx1* transcription. This interpretation is consistent with restricted spatial expression of *hesC* in *H. erythrogramma* (Extended Data Fig. 4a-d) that would seem to preclude a broad repressive function for HesC outside of the skeletogenic lineage. However, these experiments cannot rule out co-option of additional developmental functions by *hesC* in *H. erythrogramma* as the assay presented in this study only measured this transcription factor’s effect on skeletal mesenchyme differentiation. Future work aimed at validating the loss-of-function phenotype will confirm whether the function of *hesC* is completely lost or acquired novel regulatory roles during *H. erythrogramma* development. Still, altered expression of *hesC* and lack of an overt knock-down phenotype hint at a more profound evolution change within the dGRN.

We therefore turned to Pmar1, another transcriptional repressor that interacts with *hesC* to form a double-negative logic gate within the dGRN^39^: throughout most of the embryo HesC directly represses transcription of *alx1* and other genes encoding positive regulators of the skeletogenic cell fate, permitting differentiation of other cell types; in the vegetal-most cells of the embryo, however, *pmar1* is transiently expressed beginning the 16-cell stage where it represses *hesC*, allowing *alx1* transcription and thus specification of the skeletogenic cell fate^40^ (Fig.4a).

Pmar1 is encoded by a cluster of tandem genes in sea urchins^41^. We identified 10 and 20 closely linked *pmar1* paralogues in *L. variegatus* and *H. tuberculata* respectively (Supplementary Table 2). The homeodomain, nuclear localization signal, and two EH1 protein:protein interaction domains are typically well conserved, although a few likely pseudogenes are present in each species (Fig.4c, Supplementary Fig. 4). In *H. erythrogramma* we identified 11 *pmar1* paralogues (Supplementary Table 2). Surprisingly, all of these copies contain numerous substitutions, deletions, and/or frameshifts, in many cases altering or eliminating over half of the residues within the homeodomain (Fig.4c), and their expression is barely detectable at the 16-cell and 32-cell stages (Supplementary Fig. 5). In contrast, likely functional orthologs in the other two species differ by 0-3 amino acids out of 60 within the homeodomain. Furthermore, pairwise similarity between *pmar1* orthologs within a species averages greater than 88% for the entire peptide and 93% for the homeodomain in the ancestral state, while *H. erythrogramma* averages just 71.0 % and 45.3%, respectively (Fig.4d). These sequence comparisons indicate that the integrity of the *pmar1* gene family has dramatically decayed in *H. erythrogramma*, raising the question whether these genes with a crucial role in early embryonic patterning have maintained their function in the derived developmental mode.

**Figure 4.**
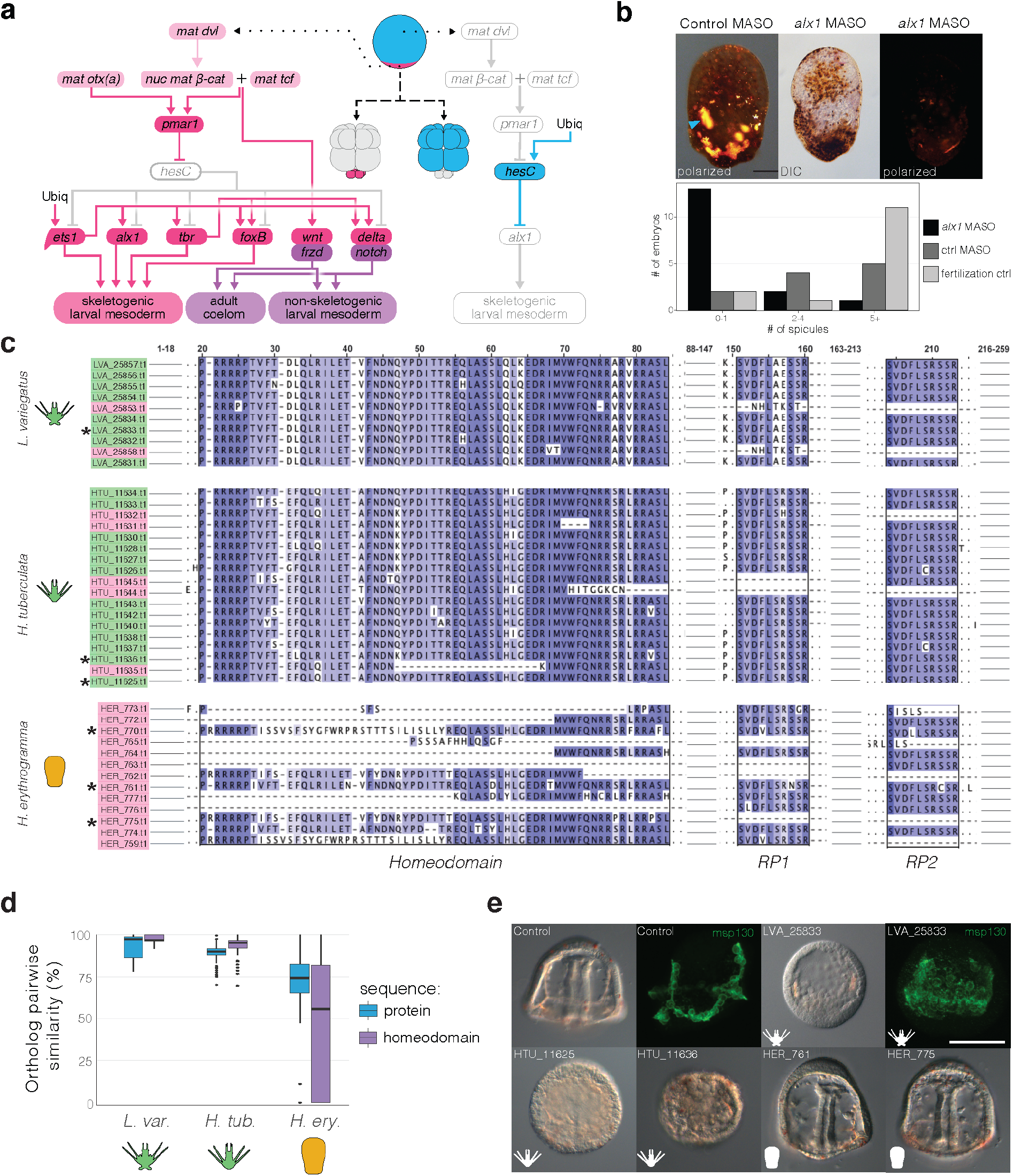
Evolutionary change at the “top” of a conserved developmental dGRN. **a**, Schematic of the ancestral dGRN that specifies skeletogenic cell fate: HesC suppresses this fate in most of the embryo (blue) but Pmar1 suppresses *hesC* in the precursors of the skeletogenic cells (magenta), where Alx1 then activates a differentiation program. **b**, Images of control and MASO knock-down of Alx1 in *H. erythrogramma* (early larva; scale bar 100µm). Polarized light (PL) illuminates skeletal elements (most are anlage of the adult skeleton, with a longer larval “arm” element out of focus on the left [blue arrow]). Bar chart of *alx1* injection summary statistics in *H. erythrogramma* (see also Supplementary Table 1). Knockdown of *alx1* expression eliminates skeleton formation in *H. erythrogramma*, as in the ancestral dGRN^36^. **c**, Alignment of homeodomain (DNA-binding) and RP domains (protein:protein interaction) from *pmar1* paralogues. Green = likely functional copy, red = predicted non-functional copy, asterisks = paralogues whose function was experimentally validated. **d**, Within-species pairwise sequence similarity of *pmar1* paralogues. Note rapid sequence divergence among paralogues in *H. erythrogramma*, and particularly within the homeodomain. Center line = median pairwise similarity; dots = outliers **e**, Overexpression assays of control and *pmar1* mRNA (prism stage; skeletogenic cells labeled with antibody that recognizes cell surface protein MSP130). DIC and fluorescent images demonstrate that mRNA of *pmar1* paralogues from *L. variegatus* and *H. tuberculata* convert most of the embryo to skeletogenic cells, whereas even the most intact *H. erythrogramma* paralogues show no such phenotype, indicating loss of function (see Supplementary Fig. 7 for additional antibody staining). dGRN: developmental gene regulatory network; MASO: morpholino antisense oligonucleotide; DIC: differential interference contrast; RP: repeated peptides.

Previous studies demonstrate that microinjecting *pmar1* mRNA into eggs produces a dramatic phenotype, with the resulting widespread overexpression of Pmar1 protein converting most of the embryo to skeletogenic cells^39,42^. Here, we utilized this assay to test the repressive function of specific *pmar1* paralogues. We separately microinjected into *L. variegatus* embryos mRNA encoding one *pmar1* paralog from *L. variegatus* and two from *H. tuberculata*. As expected, these treatments replicated the published phenotype, inducing extensive conversion to the skeletogenic cell fate, confirmed by widespread expression of the larval spicule matrix protein MSP130 (Fig.4e; Supplementary Fig. 7; Supplementary Table 3). We then separately tested the three most intact paralogues of *pmar1* from *H. erythrogramma*. At the same and higher concentrations, none was able to produce the specific or any other discernible phenotype (Fig.4e, Supplementary Fig. 6-7; Supplementary Table 3). These results indicate that the repressive role of Pmar1 is retained in *H. tuberculata* but has been lost in *H. erythrogramma*.

Together, these perturbation experiments and sequence comparisons indicate that both components of the double negative gate near the very top of the dGRN that specifies the skeletogenic cell fate do not function in *H. erythrogramma* as they do in species with the ancestral life history. Remarkably, this excision of a critical early regulatory interaction does not abort either the specification or subsequent function of skeletogenic cells: the role of Alx1, the component of the skeletogenic subcircuit immediately following the double-negative gate, remains intact (Fig.4b), structural genes characteristic of differentiated skeletogenic cells are transcribed^19^, and a simplified larval skeleton is synthesized^38^. This finding is all the more remarkable given that many other transcription factors appear to have conserved roles in *H. erythrogramma*, based on similar expression profiles and in some cases, experimental perturbation^19,23,43,44^. Taken together, these results reveal *H. erythrogramma* embryos as a mosaic of conserved and altered features that have evolved as a product of its derived life history and associated changes in selective regimes.

## DISCUSSION

Prior work showed that the evolution of nonfeeding development in *Heliocidaris* was accompanied by overt changes in oogenesis, cleavage geometry, morphogenesis, and larval morphology, with extensive underlying changes in gene expression^17-20,23^. Whole genome sequence analysis presented here demonstrates that these changes are not merely superficial consequences of amplified maternal provisioning. Although we find evidence for adaptive changes within some coding regions, these are dwarfed by the sheer number and widespread distribution of apparently adaptive changes in the sequences of putative regulatory elements and in the regulation of their chromatin states during early development (Fig.1c-g; 2). Both types of molecular change are strikingly enriched on the branch where nonfeeding development evolved and are over-represented among differentially expressed genes and especially among dGRN genes (Fig.1f-g; 2c-d). While the potential for natural selection to influence trait evolution through changes in gene regulation by altering regulatory element sequence and chromatin accessibility is widely appreciated, we are aware of few cases that illustrate the influence of both so extensively at a genomic scale and during such a short interval.

Focusing on transcriptional regulation that patterns the early embryo provides a test of the idea that evolutionary conservation of early development is the product of intrinsic constraints. We examined the earliest zygotic patterning event in the sea urchin embryo, where three transcription factors interact to specify two distinct cell fates and simultaneously establish the primary signaling center of the embryo. There is arguably no set of interactions within the dGRN that is more fundamental to patterning the early sea urchin embryo, and they are conserved among sea urchins that diverged ∼225 my ago^5^. Remarkably, however, Pmar1 and HesC, which interact to form a crucial double-negative logic gate^39^, have lost their early patterning roles in *H. erythrogramma* (Fig.4b-e, Extended Data Fig. 4). The case of *pmar1* is particularly striking, as it is present as a tandem array of genes; uniquely in the genome of *H. erythrogramma*, numerous deletions and point mutations alter about half of the homeodomain in each of 10 the paralogues, rendering their proteins nonfunctional (Fig.4c-e, Supplementary Fig.7).

The magnitude and extent of modifications to the earliest regulatory interactions within the sea urchin dGRN in *H. erythrogramma* demonstrate that some deeply conserved embryonic patterning mechanisms remain evolvable during substantial shifts in selective regimes. More broadly, conservation of gene network architecture does not necessarily imply developmental constraint, but may instead reflect long-term stabilizing selection for performance relative to a particular environment or life history. Abrupt shifts in natural selection provide valuable natural “perturbation experiments” that can reveal in detail how evolutionary mechanisms shape conservation and change in gene regulation and dGRNs organization across the tree of life.

## METHODS

### 1. Genome Sequencing and Assembly

#### 1.1 Tissue Collection

*Heliocidaris erythrogramma* (*He*) and *H. tuberculata* (*Ht*) specimens were collected near Sydney Harbor in Sydney, NSW, Australia and housed in natural sea water at the Sydney Institute of Marine Science in Mosman, NSW, AU. The interpyramidal muscle of Aristotle’s lantern (the sea urchin’s feeding apparatus), tube feet, and the ovarian tissue were dissected from a single female individual, flash-frozen in liquid nitrogen, and stored at -80C until DNA extraction and sequencing.

#### 1.2 Genomic DNA Sequencing

For each species, a 3rd-generation DNA library was sequenced on a PacBio sequel II CLR platform, generating 90.01 (*He*) and 89.47 (*Ht*) Gb of data with an N50 read length of

17.24 (*He*) and 23.70 (*Ht*) Kb. DNA from the same individual for each species was also used to construct 10x Genomics linked-reads and Hi-C libraries, which were sequenced on a BGI-SEQ 500 platform, generating 194.11 (*He*) and 199.11 (*Ht*) Gb and 130.85 (*He*) and 229.03 Gb (*Ht*) of data, respectively. Jellyfish v2.2.6^45^ and GenomeScope v1.0.0^46^ were deployed to conduct a k-mer based survey of genome composition using linked-read sequencing data based on 17-mer frequency distribution to estimate the genome size and heterozygosity of both *He* and *Ht* (Figure S4 a-b).

#### 1.3 Genome Assembly

PacBio sequencing data was employed to assemble a *de novo* contig-level genome assembly using Canu v1.8 (minReadLength=1200; minOverlapLength=1000)^47^. Subsequently, HaploMerger2 v3.6^48^ was used to create breakpoints in the contigs where potential misjoins have occurred by aligning allelic contigs via Lastz v1.02.00^49^. From these fragmented contigs, the longest of each allelic pair was identified and selected using Redundans v0.14a^50^ resulting in a near-haploid level genome assembly. The output of this pipeline was polished using Pilon v1.23^51^ with 10X sequencing data to improve assembly quality and accuracy at single base resolution. Lastly, contigs were assembled into scaffolds by mapping Hi-C read pairs to the polished assembly with HiC-Pro^52^, resulting in approximately 21.95% (*He*) and 32.10% (*Ht*) valid Hi-C reads pairs. Juicer v1.5^53^ and 3D-DNA v180419^54^ were used to correct and finalize the construction of chromosome-length scaffolds for each species. See Supplementary Fig. 8 for HiC contact maps.

#### 1.4 Repeat Identification and Classification

Genomic repetitive elements were identified with RepeatModeler v2.01^55^ to generate species-specific repeat element libraries for each. Repeat families were filtered via BLASTn v2.3.0^56^ for significant hits to gene models of the well-studied sea urchin *Strongylocentrotus purpuratus* (www.echinobase.org) to prevent unintentional masking of genic regions. Repeats were masked from the genome of each species with RepeatMasker v4.1.1 using the most sensitive setting (-s) to identify the location of repetitive elements. Long-terminal repeats were also secondarily identified with LTR_Finder v1.0.7. Outputs of both RepeatMasker and LTR_Finder were then input into RepeatCraft v1.0^57^ under default parameters to improve repeat element annotation and identification, resulting in a final genome annotation of repetitive elements. Lastly, repeats were broadly classified into functional categories described by their mode of transposition using TEclass^58^.

#### 1.5 Gene Annotation and Prediction Strategy

Previously published paired-end RNA-seq reads from six developmental stages for each *Heliocidaris* species^19^ were trimmed using Trimmomatic v0.39^59^ (TruSeq3-PE.fa: 2:30:10; leading: 3; trailing: 3; slidingwindow: 4:15; minlen: 36) and properly paired reads were mapped to their respective genomes using STAR v2.7.2. For each species, these RNA-seq alignments as well as protein models of the *S. purpuratus* v5.0 genome^60^ were input into BRAKER2^61^ (--etpmode). This program utilizes a number of additional software as a part of its pipeline including Augustus^62^, Genmark-EP+^63^, Genemark-ET^64^, DIAMOND^65^, and Samtools^66^. Gene models from the BRAKER output for filtered for transposable elements by aligning to a combined database of transposable element sequences from the MAKER gene annotations pipeline^67^ and the Dfam v3.3 transposable element database^68^ using BLAST-P^56^. Lastly, gene models were improved using the PASA pipeline^69^ by supplementing pre-existing gene models with a de-novo transcriptome retrieved from ref. 19. These gene models were annotated by aligning peptide sequences to three separate databases using BLAST-P v2.3.0^56^ : 1) *S. purpuratus* v4.2 gene models; 2) UniProt KnowledgeBase SwissProt protein models^70^; 3) RefSeq invertebrate protein models with *S. purpuratus* excluded (e-value cutoff: 1e-5)^71^ (Supplementary Data 8). The list of sea urchin gene regulatory network (GRN) genes are provided in Supplementary Data 1, retrieved from the Institute of Systems Biology (www.biotapestry.org: accessed June 27, 2017).

### 2. Whole Genome Alignment

Prior to whole genome alignment, each genome was soft-masked for repetitive elements using each species repeat element library. An optimal scoring matrix for whole genome alignment between each set of species was inferred using the *lastz_D_Wrapper.pl* script of HaploMerger2 v3.6^48^. Next whole genome alignment between each species pair was performed in both directions following UCSC guidelines outlined in the *runLastzChain.sh* and *doBlastzChainNet.pl* (https://github.com/ucscGenomeBrowser/kent) to produce .psl, .lav, .chain, and finally liftOver files for each whole genome alignment. In addition, .maf files were generated for *H. erythrogramma, H. tuberculata*, and *L. variegatus* for each chromosome using *H. erythrogramma* as the reference genome using *Multiz* and *TBA*^72^. *L. variegatus* was chosen as an outgroup species in this study because its genome was assembled and annotated using an identical sequencing and bioinformatic strategy^21^ as the two *Heliocidaris* species presented here, thereby minimizing technical bias in this regard.

### 3. ATAC-seq

#### 3.1 Sample Preparation

For each sea urchin species (*H. erythrogramma, H. tuberculata*, and *Lytechinus variegatus*), adult animals were induced to spawn via injection of 0.5 M KCl solution into the coelom. For each species, three unique male-female pairs were crossed to produce three biologically-independent replicates of sea urchin embryos. Each culture was reared in large glass dishes supplied with 20 mm filtered sea water (FSW) that was changed every six hours. Because these species exhibit different developmental rates, a conspicuous developmental milestone, shedding of the fertilization envelope at hatched blastula-stage, was selected to maximize developmental synchrony within cultures and across species for comparison. Once a culture reached the blastula stage, live embryos were collected and processed immediately for nuclei preparation and transposase treatment as a part of the ATAC-seq protocol.

#### 3.2 ATAC-seq Protocol and Sequencing

ATAC sample preparation was carried out according to the Omni-ATAC-seq protocol^73^. For each replicate, embryos were washed once in 1 mm FSW, lysed, and 50,000 nuclei were isolated for the transposition reaction as described in the Omni-ATAC-seq protocol using the Illumina TDE1 enzyme and tagmentation (TD) buffer (Cat. No. 20034197 and 20034198) (San Diego, CA, USA). Sequencing libraries for each replicate were generated via qPCR and sequencing libraries were purified and size selected using Ampure XP Beads at a 1.8:1 bead volume: library volume (Beckman Coulter, Brea, CA, USA). Library quality and transposition efficiency was accessed using a Fragment Analyzer and PROSize 2.0 (Agilent). *H. erythrogramma* and *L. variegatus* libraries were sequenced on an Illumina HiSeq 4000 instrument using 50 bp SE sequencing at an average of 41.9 million and 37.3 million reads per sample, respectively. *H. tuberculata* libraries were sequenced on an Illumina NovaSeq 6000 instrument using 50 bp PE sequencing (only SE were used for data analysis) at an average of 31.4 million reads per sample.

#### 3.3 ATAC-seq Data Analysis

Raw ATAC-seq reads were trimmed for quality and sequencing adapters using cutadapt^74^ v2.3 with the following parameters: -a CTGTCTCTTATACACATCT -q 20 --trim-n -m 40. Trimmed reads were then aligned to each species’ respective genome using stampy^75^ v.1.0.28 using the “--sensitive” set of parameters. ATAC-seq alignments were filtered for mitochondrial sequences and required an alignment quality score of at least 5 using samtools v1.9^66^.

In this study, we aimed to compare the evolution of orthologous non-coding sites. To accomplish this, we performed a series of liftOvers^76^ to convert ATAC-seq alignments between genomic coordinates of each sea urchin species (see previous section for description of genome alignments). We took an iterative, reciprocal liftOver strategy described below to minimize possible reference bias associated with converting between genome assemblies: 1) *H. erythrogramma*: *He → Lv → He*; 2) *H. tuberculata*: *Ht* → *Lv → He*; 3) *L. variegatus*: *Lv → He Lv → He*. After filtering and coordinate conversion, all ATAC-seq alignments were referenced to the *H. erythrogramma* genome with an average of 5.9 million alignments per sample to orthologous genomic loci.

Following filtering and coordinate conversion, peaks were called from these alignments using the MACS2 v2.1.2^77^ callpeak function (parameters: –nomodel, –keep-dup=auto, –shift 100, –extsize 200) for each species separately. Peak coordinates were merged using the bedtools^78^ v2.25 *merge* function requiring a peak overlap of at least 200 bp to be merged into a single peak. Lastly, for each sample, accessibility of each peak was measured with the bedtools^78^ v2.25 *multiBamCov* function.

#### 3.4 Tests for positive selection within OCRs

In order to test for evidence of positive selection, a neutral genomic reference across all species was assembled. To do this, the genome was first masked repetitive elements, coding sequence, untranslated genic sequence, non-coding RNAs (including microRNAs, ribosomal RNAs, small nuclear RNAs, and transfer RNAs) and ATAC-seq open chromatin regions (OCRs) (see below) in the genome. The remaining, putatively neutrally evolving genome was then divided into 300 bp windows, orthologous regions retrieved from each species’ genome, and filtered using the *filtering.py* and *pruning.py* scripts of the “adaptiphy”^79^ program (https://github.com/wodanaz/adaptiPhy). Next, branch lengths of each of these neutral sites was estimated using phyloFit^80^ (--subst-mod HKY85), highly-conserved sites removed (Figure S5), relative branch lengths calculated, and sites falling within the middle 50% of relative branch lengths in the *H. erythrogramma* genome were selected as the neutral reference (88,004 sites; Supplementary Data 2, Supplementary Fig. 9).

To measure branch-specific signatures of positive selection in the non-coding genome, the adaptiPhy^79^ pipeline (https://github.com/wodanaz/adaptiPhy) for global tests of natural selection was followed. First, orthologous sequences for non-coding sites of interest were selected from each species’ genome into FASTA format. Sequences were trimmed to include only contiguous DNA sequence using the *prunning*.*py* script and filtered using the *filtering*.*py* script, requiring a minimum alignment length of 75 bases. These trimmed and filtered alignments serve as “query” sequences of tests for selection. To generate a neutral reference for comparison, ten neutral sites were randomly selected (see above) and concatenated into a single neutral reference sequence. In addition, for each OCR, tests for positive selection were repeated 10 times against a unique putatively neutral reference. For each query site replicate, substitution rates of both the query and randomly concatenated neutral reference were estimated using phyloFit^80^, and the zeta score was calculated as the ratio of the query substitution rate to the neutral reference substitution rate. In addition, p-values of likelihood ratio tests for significant levels of branch-specific positive selection were calculated with adaptiPhy^79^ pipeline using HyPhy^81^. P-values and substitution rates for all query and neutral sites were then imported in R v4.0.2 for analysis (Supplementary Data 3).

#### 3.5 ATAC-seq Peak Filtering

After accessibility and rates of selection were calculated for each ATAC-seq peak, herein referred to as an “open chromatin region” (OCR), a series of filtering and quality control metrics were carried out to ensure only high confidence and quality peaks were compared between species. These filtering steps are as follows: 1) each OCR is required to have at least 75bp of contiguous, single copy sequence (see *Section 3.4*) for accurate estimations of selection; 2) for each species, a local composition complexity (LCC)^82^ value of 1.9 or more was required for the OCR to remove repetitive or other low-complexity sequences that may generate inaccurate estimations of selection (module: biopython.org/docs/1.75/api/Bio.SeqUtils.lcc.html); 3) a CPM of 3 or more was required in at least 2 (of the 9) samples to remove OCRs with extremely low accessibility; 4) the midpoint of the OCR must lie within 25kb (in either direction) of the translational start site of a gene model; 5) the gene nearest to an OCR must be the same gene in each of the species’ genomes —in other words, for each OCR and its nearest gene in the *H. erythrogramma*, the orthologous region in the *H. tuberculata* and *L. variegatus* genome must also be closest to a gene model that is orthologous (determined by annotation) to the same gene in the *H. erythrogramma* genome. Given nearly no prior knowledge is known of the cis-regulatory landscape for these sea urchin species, these stringent filtering methods were carried out in order to maximize confidence in comparisons of non-coding sequence evolution and function. This method resulted in a final set of 27,322 high-confidence OCRs for cross-species analysis (Supplementary Data 4).

#### 3.6 ATAC-seq Statistical Analysis

Raw counts of the filtered OCRs were loaded into DESeq2^83^ v1.30 to calculate differential accessibility between sample groups. For life history strategy comparisons, *H. tuberculata* and *L. variegatus* were treated as a single group. Differentially accessible sites were classified as having a 2-fold accessibility difference between sample groups and supported by an FDR of 10%. Significant levels of positive selection were classified as having a median zeta value greater than 1.5 and supported by a median false-discovery rate less than 10% across 10 replicates for each query site. Branch-specific evidence of positive selection met these criteria for one species, but failed to meet these criteria in the other, as evidenced by a zeta score < 1.5.

### 4. Coding Selection Analyses

In order to make tests for positive selection in coding sequences analogous to non-coding sequences, only single-copy orthogroups were considered in these analyses. Single-copy orthologs between *H. erythrogramma, H. tuberculata, L. variegatus*, and *Echinometra lucunter* were identified using OrthoFinder^84^. Evidence of episodic positive selection was queried on both the *H. erythrogramma* branch and *H. tuberculata* branch under default parameters using BUSTED^22^, by specifying either branch as the “foreground” branch. P-values from these analyses are available in Supplementary Data 5. Genes with significant evidence of episodic positive selection were supported by a p-value <= 10% by the likelihood ratio test.

### 5. Bulk RNA-sequencing Analysis

Raw RNA-seq reads from blastula stage embryos of *He, Ht*, and *Lv* were retrieved from ref 19, trimmed and filtered for low quality bases and reads with Trimmomatic^59^, and aligned to each species respective genomes and gene models with STAR^85^. From these alignments, mRNA expression was estimated with Salmon^86^ and loaded to R for statistical analysis. Read counts for summed to each gene’s best match to the *S. purpuratus* v4.2 gene models to generate a common reference for expression comparisons between species as described in ref 19. Differentially expressed genes between life histories were called as having a fold-change in expression > 2 and supported by an FDR of 10% or less between *He* and both planktotrophic species (in the same direction), and not DE between *Ht* and *Lv* (Supplementary Data 6).

### 6. Single-cell RNA-Sequencing

#### 6.1 H. erythrogramma embryo culturing

Female H. erythrogramma individuals were spawned via intracoelomic injection of 0.5ml of 0.5M KCl. Unfertilized eggs were washed three times in 100um filtered natural sea water (FSW). Eggs were fertilized by 2ul of concentrated sperm in .02g Para-Amino Benzoic Acid (PABA)/100 ml FSW. Following fertilization, eggs were washed three additional times in FSW to remove residual sperm and PABA. Fertilized embryos were then cultured at 22-23 °C. At 6 hours post fertilization (hpf) embryos were sampled for microscopy and dissociation, then fixed for scRNA-seq.

#### 6.2 Embryo Dissociation and Fixation

Once embryos developed to the early blastula stage (pre-skeletogenic cell ingression), a portion of the co-culture was taken, and washed one time in Calcium-Free Artificial SeaWater (CFASW). After washing embryos with CFASW, 3ml of embryos were added to 7ml of dissociation buffer made (1.0M Glycine and 0.25mM EDTA, pH 8.0) at 4 °C, and gently rocked on a rocker for 10 minutes. Following incubation, embryos were gently triturated 15-20 times to increase disassociation, then 10ml ice cold 100% methanol was added, and cells were incubated for 10 minutes and on the rocker. Following incubation, cells were triturated again 15-20x times, and then another 30ml of ice cold 100% methanol was added to bring the suspension to a final concentration of 80% methanol. This 80% methanol suspension of cells was incubated at 4 °C for 1 hour. Following this last fixation step, cells were stored at -20 °C until library preparation.

#### 6.3 Rehydration of Single Cells, Library Preparation, and Sequencing

Cells were centrifuged at 50xg, supernatant was discarded, and fixed cells were washed twice and rehydrated in a Sigma 3x Saline Sodium Citrate (SSC) buffer (SKU SRE0068) before cell count and library preparation. Cell concentration was estimated with a hemocytometer and volume was adjusted to a final concentration of ∼300 cells/ uL. Single cell libraries were prepared using the 10x Genomics 3’ v3 gene expression kit and the 10x Chromium platform to encapsulate single cells within droplets. Library quality was verified using the Agilent 2100 Bioanalyzer. In total, ∼ 3960 cells were loaded onto the 10X instrument. For the single library preparation, 2500 cells were targeted, of which 2066 were successfully captured for sequencing. Libraries were sequenced by the Duke Genomics and Computational Biology Core facility on two NovaSeq6000 S1 flow cells with 28 × 8 × 91 bp sequencing performed.

#### 6.4 *FastQ Generation, Genome Indexing, and Quantification* of scRNA-seq

Following sequencing, we used Cellranger v3.1.0 to convert Illumina-generated BCL files to fastq files using the Cellranger “mkfastq” command. scRNA-seq data for early blastula stage embryos (pre-skeletogenic cell ingression) of L*ytechinus variegatus* were retrieved from a published scRNA-seq developmental time course of the species^34^. We then applied the “mkref” command to index the most recent Lv3.0 genome^21^ (for the *Lytechinus* data) and the *H. erythrogramma* genome assembled in this study. The “count” command was used to demultiplex and count reads mapping to the respective reference *Lv* (53.3% mapping rate) or *He* (94.1% mapping rate) genome. The “mat2csv” command was used to obtain CSV RNA count matrix files for each sample for further downstream analysis. Orthofinder v2.3.12^84^ was implemented to identify putative 1-1 orthologous gene models between *Lv* and *He* (Supplementary Data 10).

#### 6.5 scRNA computational analyses

We employed a dual strategy for comparing scRNA-seq expression between *He* and *Lv*: 1) A non-integrated analysis in which scRNA-seq from each species was quantified against its own gene models (Fig. 3a) and 2) an integrated analysis in which orthologous anchor genes were used to identify cell types with overlapping expression profiles between each species (Fig. 3d). CSV RNA count matrix files were uploaded to R and a Seurat object was generated for 1) each species separately quantified against their own gene models and 2) orthologous genes between *H. erythrogramma* and *L. variegatus* (Supplementary Data 7). Each dataset was filtered to remove low quality cells with nFeature_RNA > 200, nFeature_RNA < 5500, and nCount_RNA < 7500. To ensure differences in input cell number did not bias detection of different cell types between samples, the *Lv* dataset was separately subsampled to 2065 randomly-selected cells and clustering analyses were repeated (see Extended Data Fig. 2). Furthermore, both datasets had comparable distributions of genes per cell and UMIs per cell numbers (Supplementary Fig. 2).

For the non-integrated analyses, “SCTransform”, a regularized negative binomial regression method that stabilizes variance across samples, was applied to perform normalization and removal of technical variation^87^, while preserving biological variation. We next performed Principal Component Analysis on the SCTransformed Seurat object file of raw gene expression counts, and found the nearest neighbors using 10 PC dimensions of variable gene space. UMAP (Uniform Manifold Approximation and Projection)^88^ was applied to multi-dimensional scRNA-seq data to visualize the cells in a two-dimensional space. Finally, clustering was performed using graph-based Louvain Clustering with resolution, res=0.5, resulting in 7 clusters in *L. variegatus* and 3 clusters in *H. erythrogramma*. The clusters were putatively annotated using dGRN genes and published in situ hybridization patterns as markers (Supplementary Fig. 1, Supplementary Data 11), and ambiguous cluster identities are conservatively denoted as broad embryological territories (e.g. endomesoderm, ectoderm A-C) or as “unknown”.

To perform the integrated analyses of scRNA-seq data between species, we carried out 3 independent methods for scRNA-seq data integration to compare expression of the same orthologous genes between *H. erythrogramma* and *L. variegatus* and identify putative overlapping cell types: 1) Canonical Correlation Analysis (CCA)^89^, 2) Reciprocal Principal Component Analysis (RPCA) (satijalab.org/seurat/articles/integration_rpca.html), and 3) Harmony^90^ (Fig. 3d; Extended Data Fig. 3). For each scRNA-seq strategy (non-integrated and CCA), counts for all gene models and orthologous sets of genes are included as Supplementary Data 7, and top marker genes for each cluster are available in Supplementary Data 9.

### 7. *H. erythrogramma* microinjection and in-situ hybridization

#### 7.1 MASO design and microinjection

Morpholino antisense oligos (MASOs) were constructed to target the translation start site of *alx1* (ATCAATTCGGAGTTAAGTCTCGGCA) and *hesC* (ATCCAGATGTGTTAAGCATGGTTGC) and synthesized by Gene Tools (Philomath, OR, USA). Control morpholinos included a standard negative control morpholino recommended by the manufacturer (CCTCTTACCTCAGTTACAATTTATA) and a scrambled morpholino for HesC (ATCGACATCTGTTAACCATCGTTGC). Fertilized eggs of *H. erythrogramma* were injected as described in ref. 91 at a concentration of 100 µM and 200 µM for *alx1* and 500 µM for *hesC*, then reared at 22 °C in pasteurized, 0.22-micron filtered seawater + penicillin (100 unit/mL) and streptomycin sulfate (0.1 mg/mL) (Sigma P4333A). Injected embryos were checked for developmental abnormalities and mortality every six hours until fixation.

#### 7.2 Fixation and ISH

*H. erythrogramma* embryos were fixed for in-situ hybridization (ISH) for ∼16 hrs overnight at 4 °C in 4% paraformaldehyde (Sigma 158127) + 20 mM EPPS (Sigma E1894) in FSW, washed 3x in FSW, and dehydrated step-wise into 100% MeOH and stored at -20 °C. The full-length *HesC* coding sequence was synthesized in vitro by GenScript (Piscataway, NJ, USA) and subcloned (NCBI insert number: MK749159), and used as template to make antisense RNA probes for ISH. ISH of *H. erythrogramma* was performed according to previously published methods^44^. Hybridizations were carried out at 65 °C and stringency washed at 0.1% SSC.

#### 7.3 Imaging

Fixed *H. erythrogramma* embryos were washed with 100% EtOH, cleared and mounted in 2:1 (v/v) benzyl benzoate: benzyl alcohol (BB:BA). DIC or polarized light (PL) micrographs were taken on Olympus BX60 upright microscope with an Olympus DP73 camera. ISH images were taken on a Zeiss Upright AxioImager with a Zeiss MRm camera using ZEN Pro 2012 software.

### 8. Pmar1 mRNA overexpression assays

#### 8.1 mRNA Synthesis

Sequences of *pmar1* orthologs were retrieved from each species respective genome annotations (*Lv*^*21*^, *He* and *Ht*, this study) (Supplementary Table 2). One *Lv* ortholog (LVA_25833.t1) was selected for overexpression assays as it represents the ortholog tested in previous overexpression assays^42^, while two *Ht* (HTU_11636.t1 and HTU_11625.t1) and three *He* (HER_770.t1, HER_761.t1, and HER_775.t1) were selected for overexpression assays as they represent orthologs with the highest identity to the species’ consensus sequence, and therefore predicted as genes most likely to be functional. Construct templates for each ortholog were ordered from Twist Biosciences (San Francisco, CA, USA) and mRNA was synthesized from these constructs with a ThermoFisher MEGAshortscript T7 Transcript Kit (AM1354).

#### 8.2 Overexpression experiments

Female *L. variegatus* individuals were spawned via intracoelomic injection of 0.5M KCl, washed in FSW, and fertilized with 1 uL of concentrated sperm in 100 ml FSW. For each construct, at least 4 rounds of microinjections were conducted with each round including 30-50 healthy embryos. *Lv* constructs were injected at a concentration of 250ng/µL. *Ht* constructs were injected at a concentration of 1200ng/µL, and *He* constructs were injected at 1500ng/µL. Higher concentrations of *Heliocidaris* constructs were used to reproducibly obtain the skeletal cell conversion phenotype, and reduced sensitivity of these assays may be attributable to less optimal cross-species interactions of Pmar1 in regulating the *L. variegatus* genome. Following injection, embryos were incubated at 23 °C and imaged at 24 hpf.

#### 8.3 Imaging and Immunostaining

At 24 hpf, live embryos from each experiment were mounted on slides and imaged using DIC microscopy. Embryos were imaged on a Zeiss Axioplan II upright microscope controlled by Zen software. Also at 24 hpf selected embryos were fixed in 100% ice cold methanol. Immunostaining was carried out as described in ref. 92 to mark expression of Msp130 protein. Blocking and incubation of the secondary antibody was increased to 1 hr. Incubation of the primary antibody was set to 48 hours. A Zeiss 880 inverted confocal Airyscan microscope controlled by Zen software was used to take Z-stack images of stained embryos. Pmar1 overexpression results are summarized in Supplementary Table 3.

### 9. Data Availability

#### 9.1 Genomes

Sequencing reads used to assemble the *Heliocidaris* genomes and the genome assemblies themselves are available on the Chinese National GeneBank (CNP0002233). Genome assemblies of *Heliocidaris erythrogramma* and *Heliocidaris tuberculata* are also available on NCBI (PRJNA827916 and PRJNA827769, respectively). Genome assembly of *Lytechinus variegatus* is previously published^21^ and available on NCBI (PRJNA657258). Genome annotations and files associated with whole genome alignments between species are available on Dryad (https://doi.org/10.5061/dryad.sj3tx966v).

#### 9.2 ATAC-seq

Raw sequencing reads for the ATAC-seq dataset are available on NCBI (PRJNA828607). Alignment files are available on Dryad (https://doi.org/10.5061/dryad.sj3tx966v). Result files associated with the ATAC-seq analyses are available as Supplementary Data 2-4.

#### 9.3 Bulk RNA-seq

Bulk RNA-seq data was retrieved from Israel et al, 2016^19^.

#### 9.4 Single cell RNA-seq

Raw sequencing reads for the *Heliocidaris erythrogramma* single cell ATAC-seq dataset are available on NCBI (PRJNA833141). Sequencing reads from the *Lytechinus variegatus* single cell RNA-seq dataset were retrieved from Massri et al, 2021^34^ and available on NCBI (PRJNA765003). Results files associated with the single cell RNA-seq analyses are available as Supplementary Data 7,9.

## AUTHOR CONTRIBUTIONS

LW, MB, GF, and GAW conceived and designed the study. LW, DK, and PC collected tissues for genomic sequencing. YZ performed genomic DNA extraction and sequencing library preparations. PLD, HG, and HZ assembled the genomes. PLD performed genome annotation, alignments, and data analysis. PLD collected, prepared, and analyzed ATAC-seq libraries. AJM collected, prepared, and analyzed single cell RNA-seq libraries. JSS and AE performed embryonic expression and injection assays. PLD and GAW wrote the manuscript and all authors contributed to manuscript revisions.

## ACKNOWLEDGEMENTS

We thank the Sydney Institute of Marine Science (SIMS) for facilities, as well as the SIMS staff for their assistance. This work was supported by the National Science Foundation Division of Integrative Organismal Systems (award no. 1929934 to GAW) and National Science Foundation Graduate Research Fellowships to HRD and AJM.

## MATERIALS & CORRESPONDENCE

Correspondence and material requests regarding this work should be addressed to Dr. Gregory Wray at.

## EXTENDED DATA FIGURE LEGENDS

**Extended Data Figure 1.**
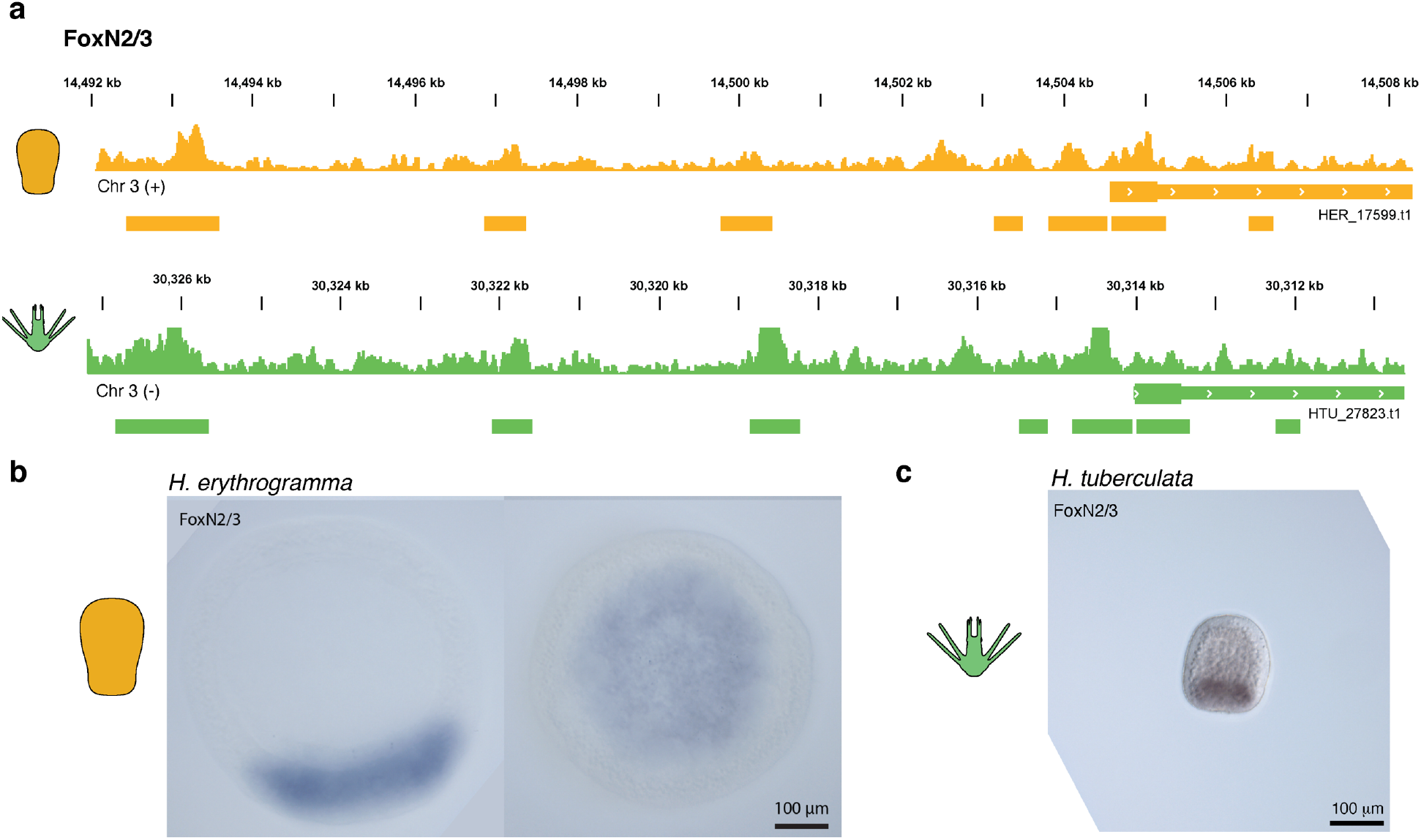
Chromatin landscape and expression domain of *foxN2/3* is conserved. **a**, Chromatin accessibility nearby *foxN2/3* in *H. erythrogramma* (top, orange) and *H. tuberculata* (bottom, green), including seven open chromatin regions (OCRs). *In-situ* hybridization of *foxN2/3* in blastula-stage **b**, *H. erythrogramma* and **c**, *H. tuberculata* embryos.

**Extended Data Figure 2:**
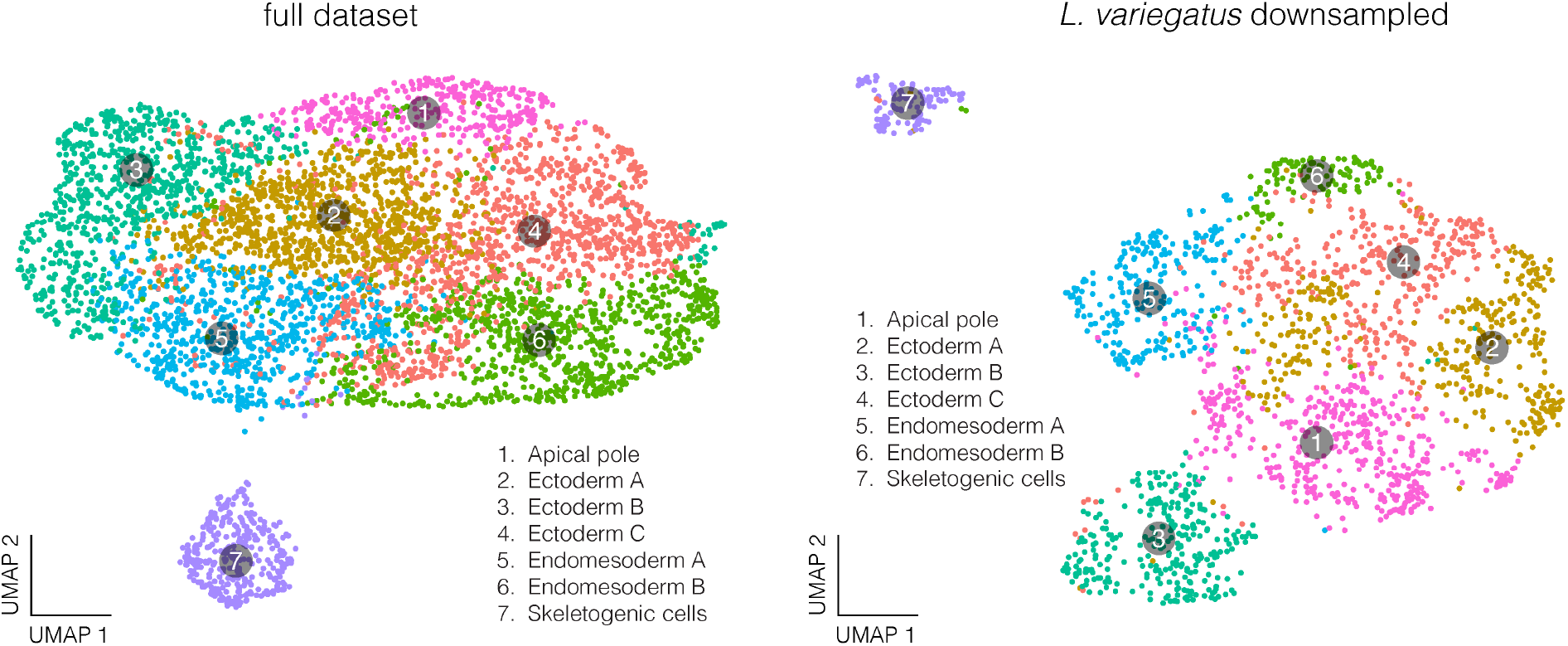
Scaling down number of *L. variegatus* cells does not significantly affect clustering. UMAP of non-integrated single cell RNA-seq data from *L. variegatus*, in which the data has been randomly subsampled to 2065 cells so that the cell number is equivalent to the *H. erythrogramma* dataset. This subsampling does not change the number and general spatial relationship among clusters in *L. variegatus* (i.e. distinct cluster of skeletogenic cells separate from remainder of cells in the embryo).

**Extended Data Figure 3:**
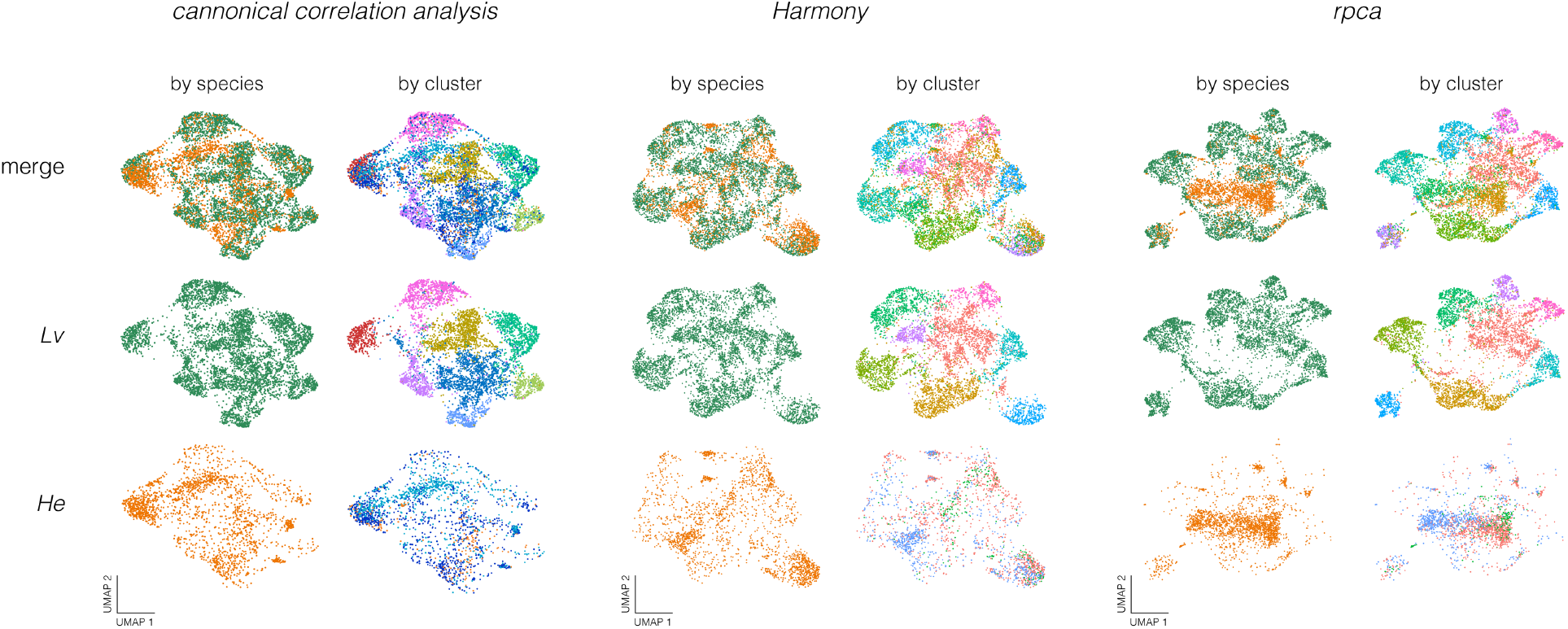
Three independent methods of integrating single cell RNA-seq data recover similar clustering relationships among *L. variegatus* and *H. erythrogramma* cells. Canonical correlation analysis (CCA), reciprocal principal component analysis (RPCA), and Harmony each recover more numerous, distinct transcriptional states in *L. variegatus* relative to *H. erythrogramma* following integration of single cell expression data. All three methods reveal a consistent number of clusters in both species. In particular, *H. erythrogramma* contains fewer clusters there is more extensive overlap of cells among clusters. These results suggesting that differentiation of discrete cell populations is delayed in the *H. erythrogramma* embryo.

**Extended Data Figure 4:**
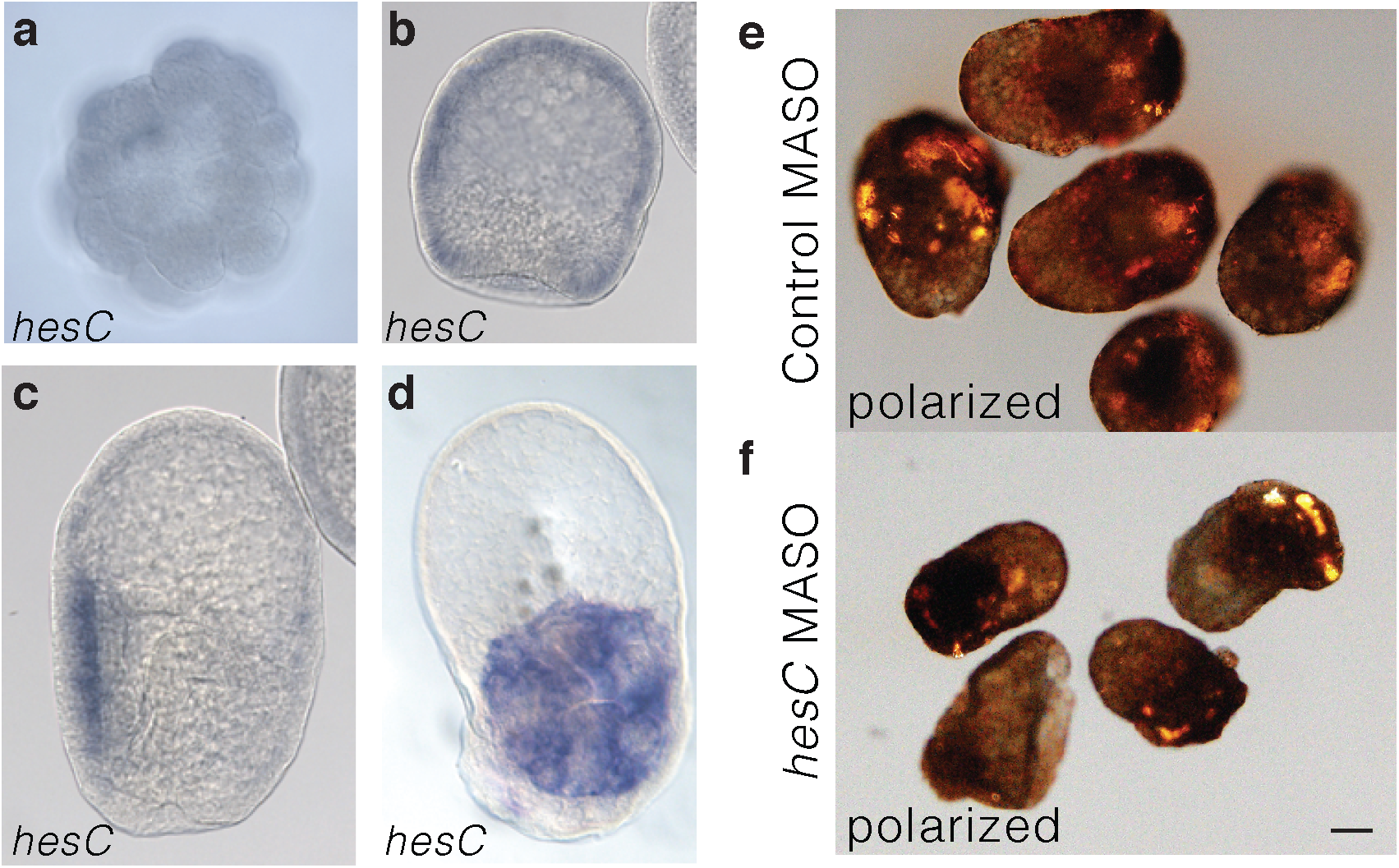
*hesC* appears to have lost its ancestral role of repressing larval skeletal cell specification in *H. erythrogramma*. Derived expression patterns of *hesC* in *H. erythrogramma* at **a**, cleavage; **b**, blastula; **and c-d**, larva stage embryos. **e-f**, Control and MASO knock-down of HesC in *H. erythrogramma* (early larva; scale bar 100µm). Polarized light illuminates skeletal elements. HesC knockdown appears to show no phenotype, a dramatic change from the ancestral dGRN.

