## Supplementary Figures and Tables for "Recent reconfiguration of an ancient developmental gene regulatory network in *Heliocidaris* sea urchins"

for

| **Date** | **Individual** | **Treatment** | **Total Spicules** |
|---|---|---|---|
| *2017.01.24* | F1 | fert.ctrl | 5 |
| *2017.01.24* | F2 | fert.ctrl | numerous |
| *2017.01.24* | F3 | fert.ctrl | 4 |
| *2017.01.24* | F4 | fert.ctrl | 1 |
| *2017.01.24* | F5 | fert.ctrl | numerous |
| *2017.01.24* | F6 | fert.ctrl | numerous |
| *2017.01.24* | F7 | fert.ctrl | numerous |
| *2017.01.24* | F8 | fert.ctrl | numerous |
| *2017.01.24* | F9 | fert.ctrl | numerous |
| *2017.01.24* | F10 | fert.ctrl | 0 |
| *2017.01.24* | A1 | alx1.mo.100uM | 0 |
| *2017.01.24* | A2 | alx1.mo.100uM | 0 |
| *2017.01.24* | A3 | alx1.mo.100uM | 0 |
| *2017.01.24* | A4 | alx1.mo.100uM | 0 |
| *2017.01.24* | A5 | alx1.mo.100uM | 4 |
| *2017.01.24* | A6 | alx1.mo.100uM | 2 |
| *2017.01.24* | A7 | alx1.mo.100uM | 1 |
| *2017.01.24* | A8 | alx1.mo.100uM | 0 |
| *2017.01.24* | A9 | alx1.mo.100uM | 1 |
| *2017.01.24* | A10 | alx1.mo.100uM | 0 |
| *2017.01.24* | A11 | alx1.mo.100uM | numerous |
| *2017.01.24* | A12 | alx1.mo.100uM | 0 |
| *2017.01.24* | C1 | std.mo.100uM | 4 |
| *2017.01.24* | C2 | std.mo.100uM | numerous |
| *2017.01.24* | C3 | std.mo.100uM | 5 |
| *2017.01.24* | C4 | std.mo.100uM | 4 |
| *2017.01.24* | C5 | std.mo.100uM | 0 |
| *2017.01.24* | C6 | std.mo.100uM | numerous |
| *2017.11.23* | A13 | alx1.mo.200uM | 0 |
| *2017.11.23* | A14 | alx1.mo.200uM | 0 |
| *2017.11.23* | A15 | alx1.mo.200uM | 0 |
| *2017.11.23* | A16 | alx1.mo.200uM | 0 |
| *2017.11.23* | C7 | std.mo.200uM | numerous |
| *2017.11.23* | C8 | std.mo.200uM | 0 |
| *2017.11.23* | C9 | std.mo.200uM | 4 |
| *2017.11.23* | C10 | std.mo.200uM | 5 |
| *2017.11.23* | C11 | std.mo.200uM | 2 |

| | | | |
|---|---|---|---|
| *2017.11.23* | F11 | fert.ctrl | numerous |
| *2017.11.23* | F12 | fert.ctrl | numerous |
| *2017.11.23* | F13 | fert.ctrl | numerous |
| *2017.11.23* | F14 | fert.ctrl | numerous |

**Supplementary Table 1**: Injection statistics for *alx1* morpholino antisense oligo injections in *H. erythrogramma*. "Numerous" denotes individuals with too many spicules to accurately quantify, typically >> 5.

| *Heliocidaris erythrogramma* | *Heliocidaris tuberculata* | *Lytechinus variegatus* |
|---|---|---|
| HER_759.t1 | **HTU_11625.t1** | LVA_25831.t1 |
| **HER_761.t1** | HTU_11626.t1 | LVA_25832.t1 |
| HER_762.t1 | HTU_11627.t1 | **LVA_25833.t1** |
| HER_763.t1 | HTU_11628.t1 | LVA_25834.t1 |
| HER_764.t1 | HTU_11630.t1 | LVA_25853.t1 |
| HER_765.t1 | HTU_11631.t1 | LVA_25854.t1 |
| **HER_770.t1** | HTU_11632.t1 | LVA_25855.t1 |
| HER_772.t1 | HTU_11633.t1_...* | LVA_25856.t1 |
| HER_773.t1 | HTU_11634.t1 | LVA_25857.t1 |
| HER_774.t1 | HTU_11635.t1 | LVA_25858.t1 |
| **HER_775.t1** | **HTU_11636.t1** | |
| HER_776.t1 | HTU_11637.t1 | |
| HER_777.t1 | HTU_11638.t1 | |
| | HTU_11640.t1 | |
| | HTU_11642.t1 | |
| | HTU_11643.t1 | |
| | HTU_11644.t1 | |
| | HTU_11645.t1 | |

**Supplementary Table 2:** Gene IDs of *pmar1* paralogues for each focal species of this study. *Complete ID: HTU_11633.t1_HTU_11639.t1_HTU_11641.t1. **Bolded** gene models are those experimentally tested in overexpression assays.

| | | | *phenotype of injected embryos (~%)* | | |
|---|---|---|---|---|---|
| **Species** | **Pmar1 construct** | **# Rounds** | **None (normal)** | **Weak (partial)** | **Strong (full)** |
| *Lv* | LVA_25833 | 8 | 10 | 40 | 50 |
| *Ht* | HTU_11636 | 7 | 20 | 30 | 50 |
| *Ht* | HTU_11625 | 6 | 60 | 20 | 20 |
| *He* | HER_770 | 4 | 100 | 0 | 0 |
| *He* | HER_761 | 4 | 100 | 0 | 0 |
| *He* | HER_775 | 4 | 100 | 0 | 0 |

**Supplementary Table 3: Summary statistics of *pmar1* overexpression injection experiments.** Each injection round typically included 30-50 healthy *Lytechinus variegatus* embryos. "# Rounds" is the number of independent rounds of microinjection carried out for each construct. "Weak" phenotype is a qualitative reference to a partial conversion of embryonic cells to primary mesenchyme cells (PMCs). "Strong" phenotype refers to a nearly complete conversion of embryonic cells to PMCs. Percentages are an approximated average across injection rounds (replicates) for each construct. *Lv*: *Lytechinus variegatus*; *Ht*: *Heliocidaris tuberculata*; *He*: *Heliocidaris erythrogramma*.

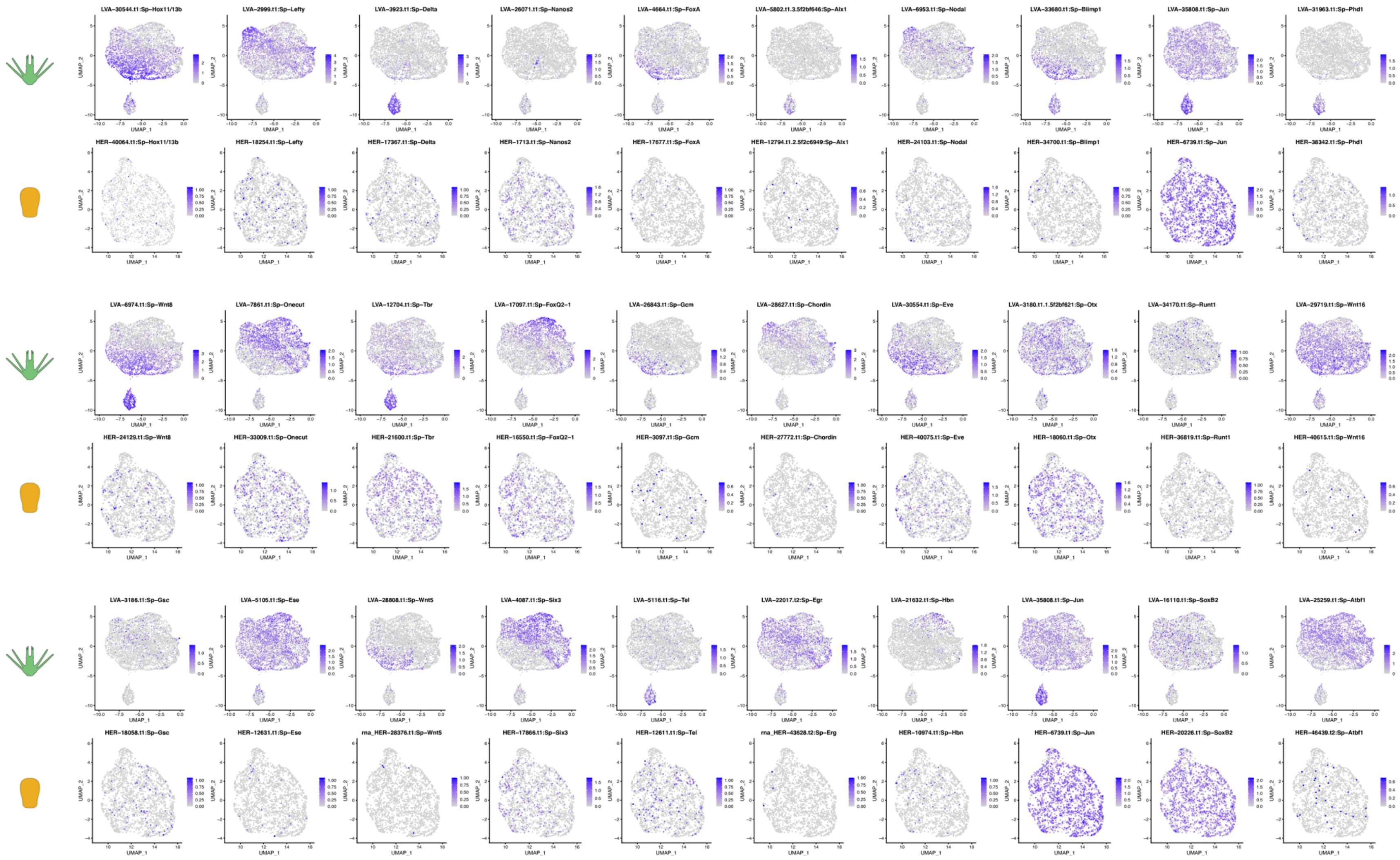

**Supplementary Figure 1. Distribution of marker genes used to determine cluster identity.** Projections show the distribution of examples of the marker genes that were used for cluster identification. See echinobase.org for more information about individual genes and references to original studies.

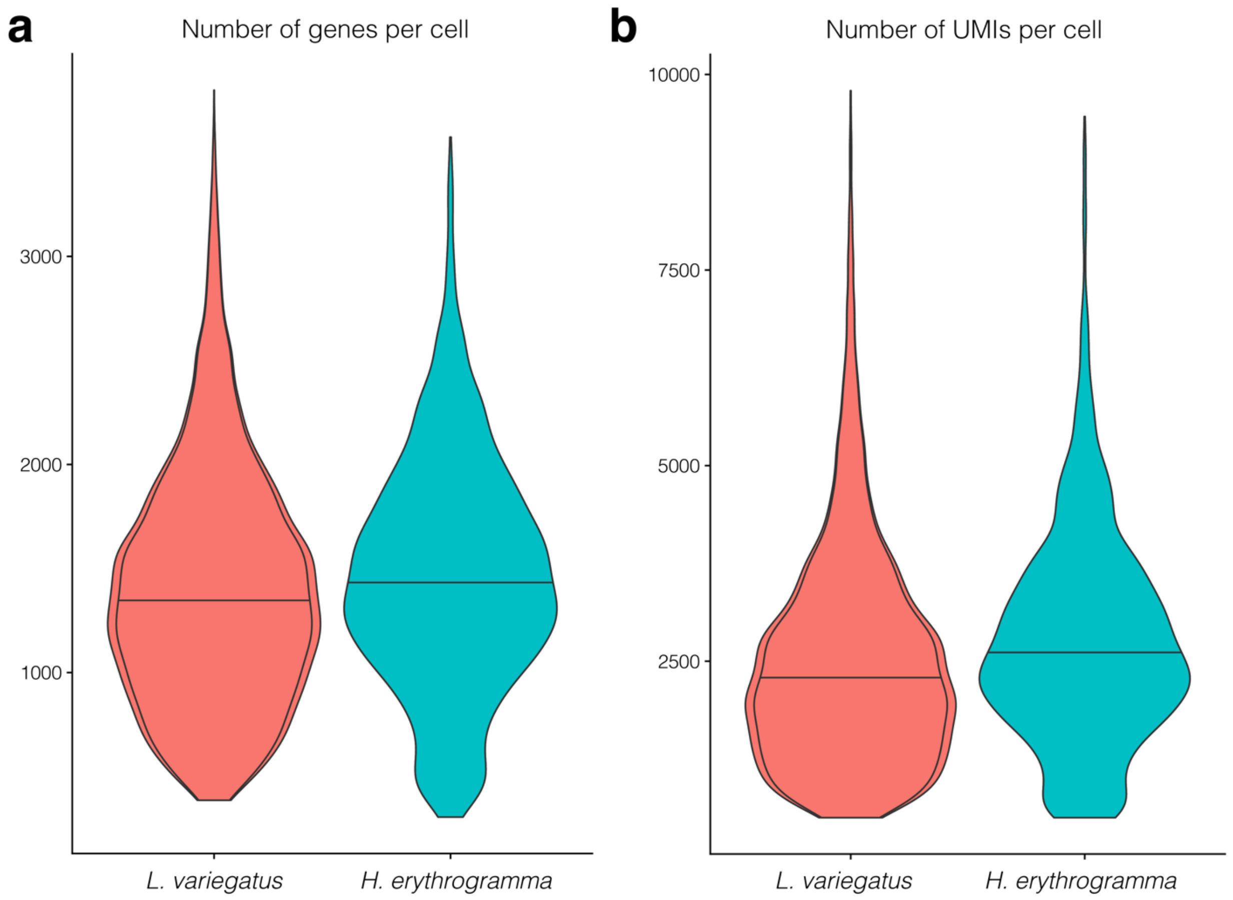

**Supplementary Figure 2:** *L. variegatus* and *H. erythrogramma* scRNA-seq libraries are of comparable quality in terms of the number of **a)** genes per cell and **b)** UMIs per cell**.**

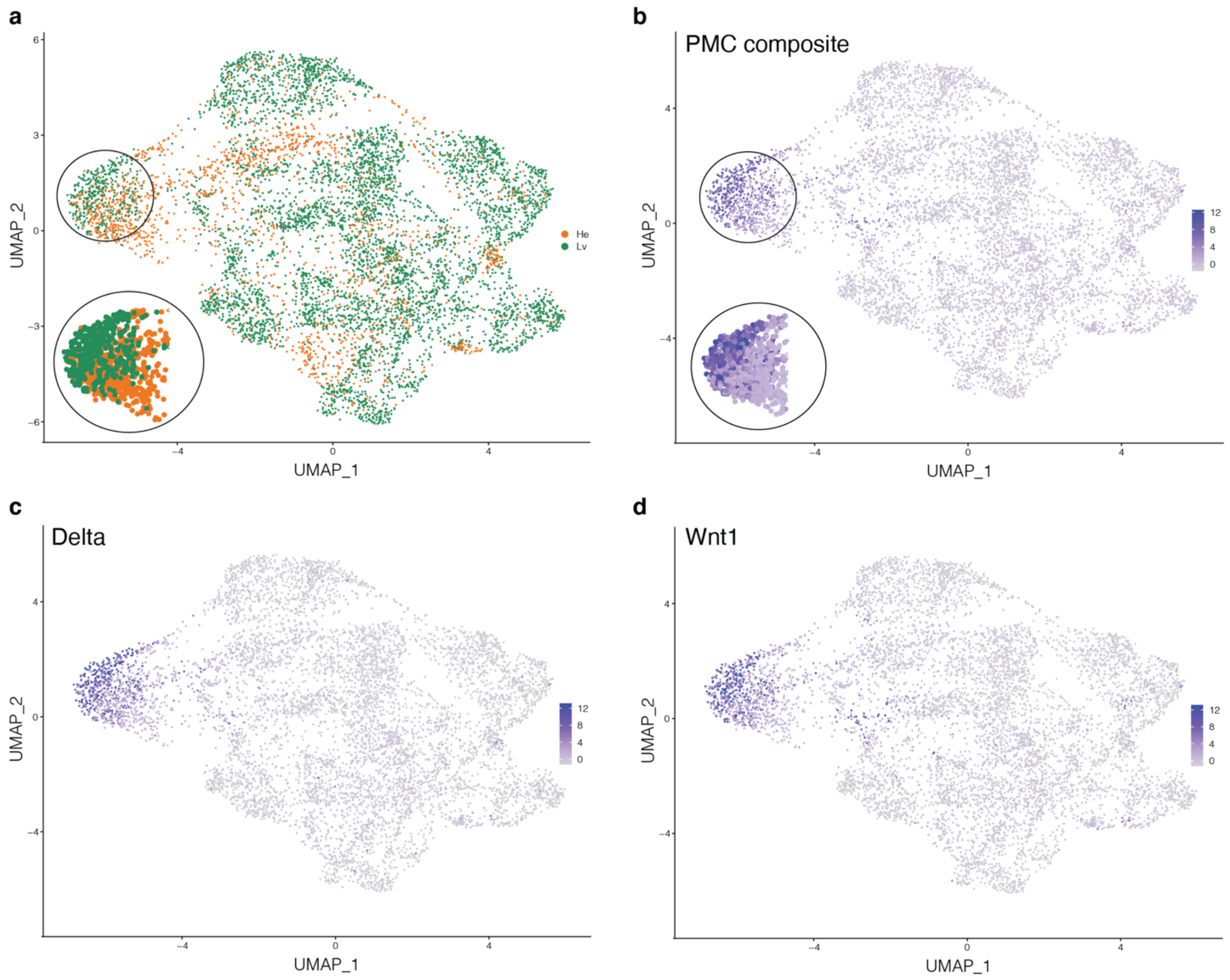

**Supplementary Figure 3: Expression markers of primary mesenchyme cells are detected in *L. variegatus*, but less so in *H. erythrogramma*. a,** UMAP of integrated single cell RNA-seq data, colored by species (from Fig. 3b). **b,** Composite expression of PMC marker genes from the same dataset, including **c,** *delta* and **d,** *wnt1*. Insets are magnified areas of the putative PMC domain, demonstrating the strongest PMC signature is found in cells belonging to *L. variegatus*.

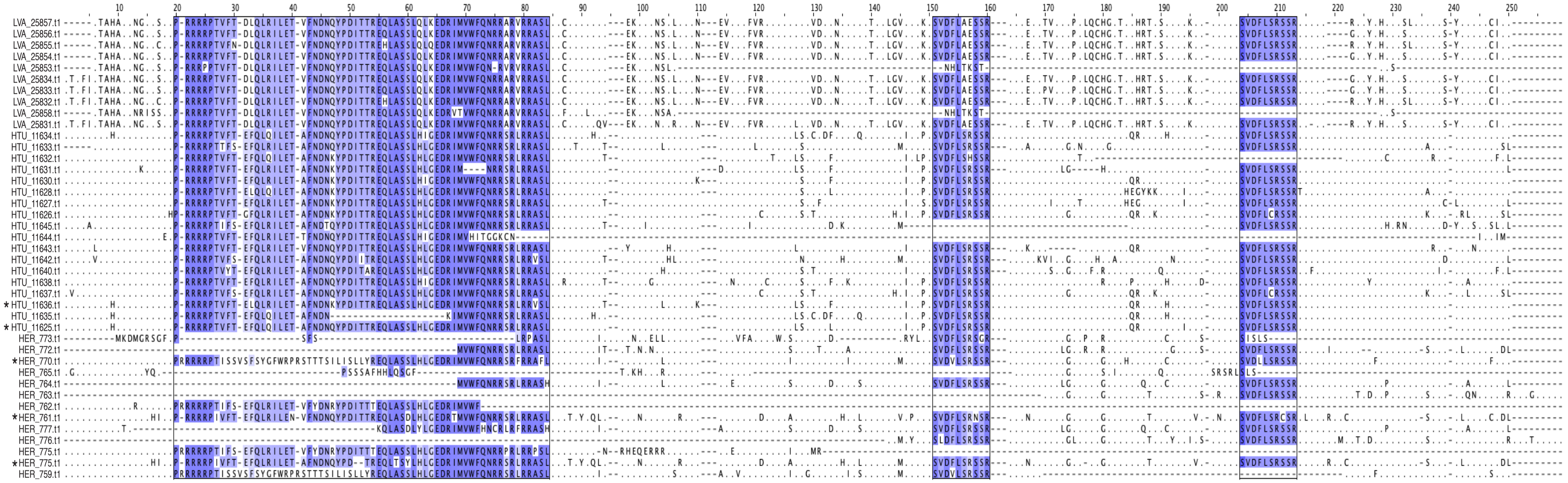

**Supplementary Figure 4:** Alignment of entire peptide sequence of each *pmar1* paralogue from *H. erythrogramma*, *H. tuberculata*, and *L. variegatus*. Homeodomain and two repeated peptide domains discussed in main text are highlighted.

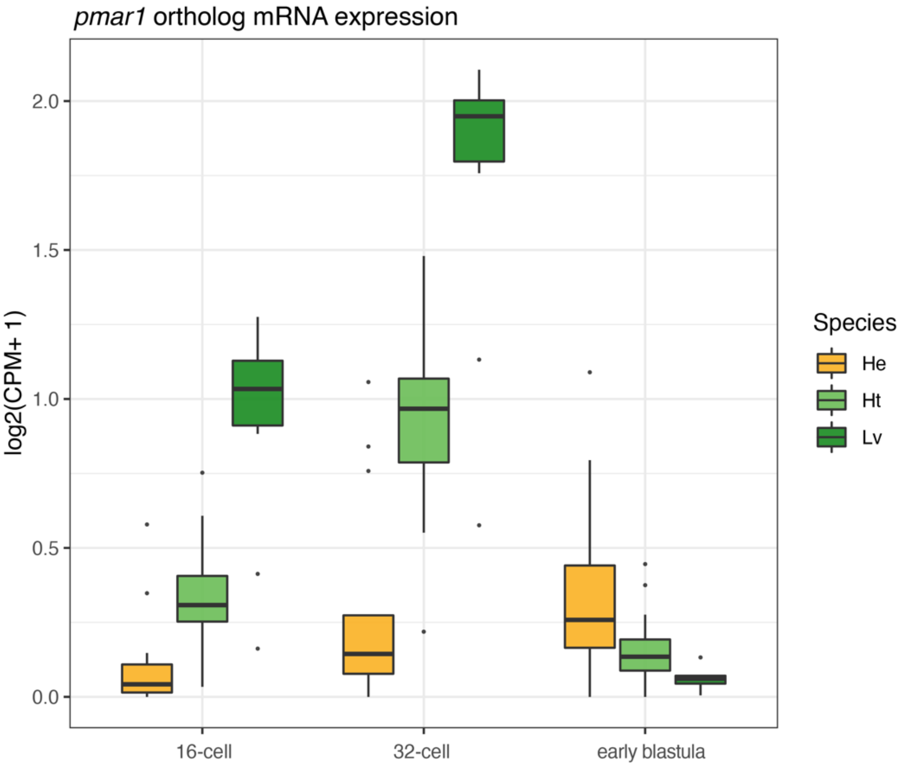

**Supplementary Figure 5:** Expression of *pmar1* paralogues from whole embryo RNA-seq[1] at 16-cell, 32-cell, and early blastula stage embryos for *H. erythrogramma* (*He*), *H. tuberculata* (*Ht*), *L. variegatus* (*Lv*).

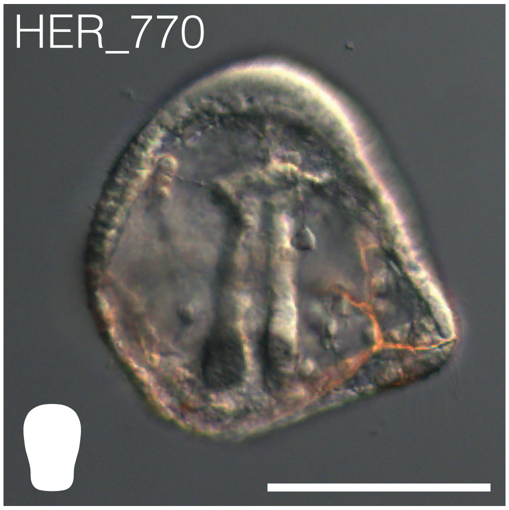

**Supplementary Figure 6:** DIC image of overexpression assay of HER_770.t1 *pmar1* paralogue in *L. variegatus*. Scale bar: 50 µm.

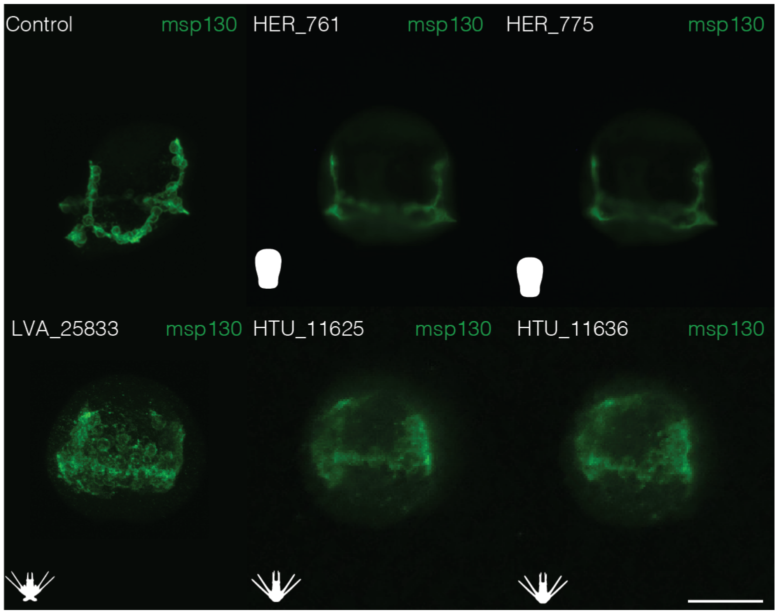

**Supplementary Figure 7:** Antibody stains for MSP130 in *L. variegatus* embryos. *Top*: Control and embryos injected with *pmar1* constructs from *H. erythrogramma* show no skeletal phenotype and develop normally. *Bottom*: Embryos injected with *pmar1* constructs from *L. variegatus* and *H. tuberculata* induce conversion of embryonic cells to primary mesenchyme cells and mesodermal defects. Control and *L. variegatus* results from Fig. 4e in the main text. Scale bar: 50 µm.

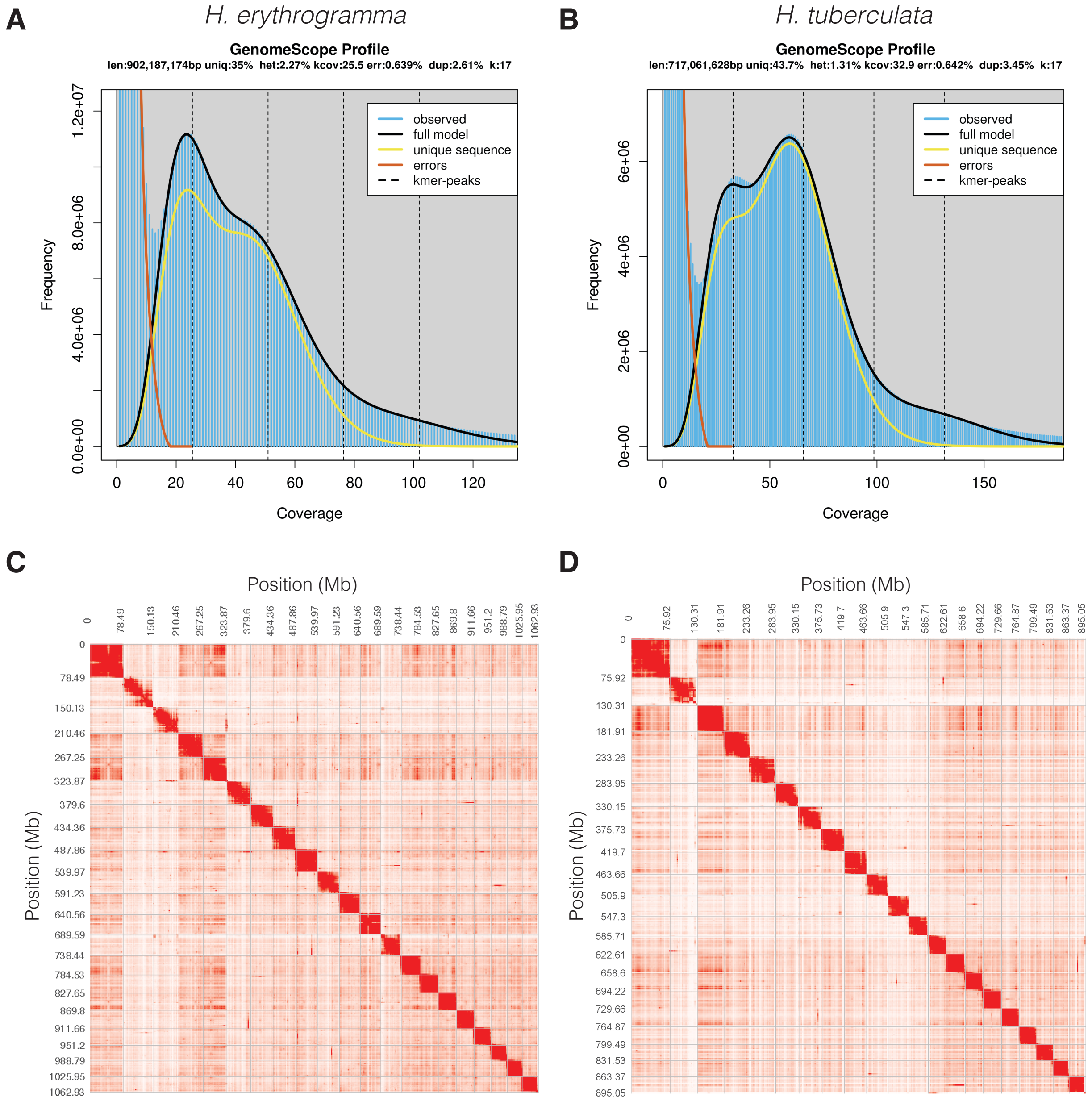

**Supplementary Figure 8:** GenomeScope profiles of **a,** *H. erythrogramma* and **b,** *H. tuberculata* genome assemblies presented in this study. Contact matrix generated from HiC data marking clear interaction boundaries delineating 21 chromosome-length scaffolds for both **c,** *H. erythrogramma* and **d,** *H. tuberculata*.

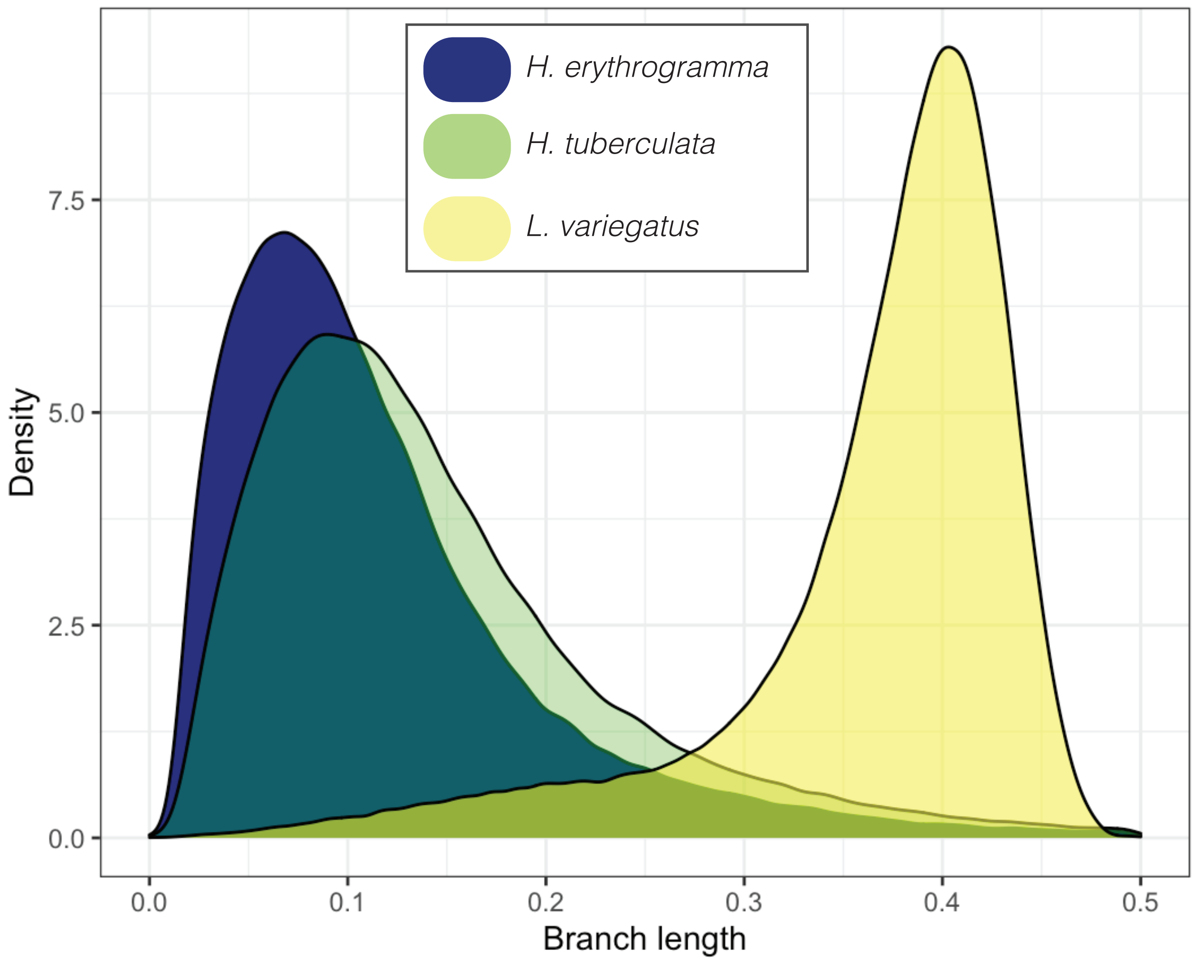

**Supplementary Figure 9:** Branch length distribution of putatively neutral non-coding sites for each sea urchin species analyzed in this study. The middle 50% of sites in the *H. erythrogramma* distribution (and corresponding orthologous sites in *H. tuberculata* and *L. variegatus*) were selected to represent the neutral reference for tests for positive selection.
